# A balance between environmental filtering and competitive exclusion modulates the macroecology of alternative stable states in microbial communities

**DOI:** 10.64898/2025.12.19.695337

**Authors:** Annamaria Bupu, William R. Shoemaker, Onofrio Mazzarisi, Jacopo Grilli

## Abstract

Predicting the assembly and stability of microbial ecosystems is fundamentally challenged by the stochasticity inherent in historical contingencies. Although these dynamics often appear reproducible at coarse taxonomic or functional scales, the mechanisms governing fine-grained species-level variation and the emergence of alternative stable states remain an unresolved problem in ecology. Bridging the gap between individual species variability and the emergence of predictable community-level attractors is essential for a mechanistic understanding of ecosystem resilience. Here, by analyzing replicate microbial communities assembled in vitro, we demonstrate that community structure is built upon individual species exhibiting robustly bimodal abundance distributions. Each taxon exists in either a high- or low-abundance state. We show that the collective configuration of these binary species states—a phenomenon also observed in natural systems—drives the correlation structure between community members and defines a set of discrete, alternative community-level attractors. By examining the prevalence of species across these attractors, we uncover a striking taxonomic signal: phylogenetically close relatives are significantly more likely to display reciprocal prevalence patterns—where the dominance of one relative necessitates the suppression of the other—than expected by chance. This discovery directly rejects models based on simple functional redundancy. Instead, it proves that state-dependent competitive exclusion is a primary driver of community divergence, where the identity of the “winner” is contingent on the collective state of the entire community. Our work reframes microbial community structure through the lens of discrete population states, providing a predictive framework for understanding ecosystem stability and the engineering of community functions.

## Introduction

Microbial communities are remarkably complex, with their composition varying significantly across space and time. A central goal of microbial ecology is to understand the ecological processes that drive this variability. A major driver shaping abundance fluctuations is the variability of the environment. Changes in biotic and abiotic factors alter species’ physiology, determining birth or death rates, and ultimately their abundance.

Environmental variability is a major source of the dynamics of natural [1–3] and experimental [4] microbial communities. Over long timescales, in the absence of major environmental shifts, it can explain fluctuations around a single, reproducible community composition. But the structure of variation can be more complex, and reflects the existence of multiple alternative stable states, or attractors, where different community compositions can persist over long times under identical environmental conditions. The existence of alternative stable states is a foundational concept in ecology [5], with solid theoretical [6] and empirical [7] ground. In microbial ecology, variation in the composition of the gut microbiome across host have been interpreted as alternative stable states [8]. Disentangling the nature of variability, uncovering the existence of alternative stable states, is crucial for predicting the outcome of community assembly and engineering desired ecosystem functions.

Experimentally assembled communities offer a powerful platform for investigating the fundamental principles of community assembly in controlled, replicable conditions. The emergence of alternative states is often a hallmark of historical contingency, where stochastic events, such as the timing and order of species arrival—so-called priority effects—can steer a community towards different and persistent compositional endpoints [9, 10]. Previous work using such systems has shown that, despite high initial diversity, communities often converge to a limited number of reproducible compositional states that are well defined by their composition at coarse taxonomic (e.g., family) levels [11, 12].

This convergence at coarse taxonomic (e.g., family) or functional compositions to a small set of states together with high-variability at finer taxonomic scales (e.g., at the level of Amplicon Sequence Variants, ASVs) is a widely discussed observation in microbial ecology, across disparate environments as well as across spatial and temporal scales [13–16]. The observation of functional convergence could be the outcome of functional selection — a particular function is environmentally selected in the community context — or a statistical artifact — a function is more reproducible than fine-scale taxonomic composition simply because it is the outcome of interactions between a large number of community members whose weakly correlated fluctuations average out. Evidence shows that functional selection is supported in some systems but not in others [17].

Such studies have established the existence of attractors at the level of taxonomic families as well as community functions [11, 12, 18]. However, the fine-grained structure of taxonomic variation around a given attractor has remained unresolved, even under identical experimental conditions. Our lack of understanding at fine-scale limits our ability to predict the outcomes of community assembly. One expects fine-grained taxonomy patterns of variation across communities to reflect both the existence of multiple attractors and the variation within a given attractor as the outcomes of functional selection [17].

Macroecological frameworks provide a powerful lens for describing microbial community structure, showing that species in natural ecosystems typically exhibit unimodal Abundance Fluctuation Distributions (AFDs), consistent with populations fluctuating around a single characteristic abundance [2, 4, 19]. In natural communities similar ASVs show positive abundance correlations as a result of environmental fluctuations [3], suggesting that limiting similarity — where competition excludes taxa with overlapping niches — plays a less important role. It is unknown whether *in vitro* communities display similar patterns of abundance correlations, given that they are assembled under controlled, reproducible, conditions. Moreover it is not known how the existence of multiple attractors, functional convergence, and the redundancy of functions across community members affect those statistical patterns.

Applying established ecological theory is crucial for moving from descriptive studies towards a predictive understanding of microbial communities [20]. While some studies in controlled microcosms suggest that assembly can follow simple, predictable rules [21], the fine-grained compositional patterns that define more complex outcomes such as alternative stable states remains poorly understood. Here, we investigate the fine-grained compositional variability at the ASV level in replicate communities assembled *in vitro* from single and multiple soil inocula. We find that, contrary to patterns in natural ecosystems, the vast majority of ASVs display a starkly bimodal AFD, corresponding to two distinct abundance states—high and low. We demonstrate that the collective switching of ASVs between these binary states defines a set of alternative community-level attractors. By analyzing the behavior of ASVs across these attractors, we find that phylogenetically close relatives are significantly more likely to exhibit reciprocal prevalence patterns, providing strong evidence for niche differentiation and competitive interactions as key drivers shaping these alternative stable states.

## Results

### ASVs have a bimodal abundance distribution

The Abundance Fluctuation Distribution (AFD) describes the fluctuations of (relative) abundances of a given ASV across communities (illustrated in Figure 1A). Natural communities of multiple systems display unimodal AFDs [2, 19, 22, 23]. A unimodal AFD implies the existence of a typical abundance that a given ASV fluctuates about (e.g., corresponding to the average abundance calculated across samples).

**Figure 1:**
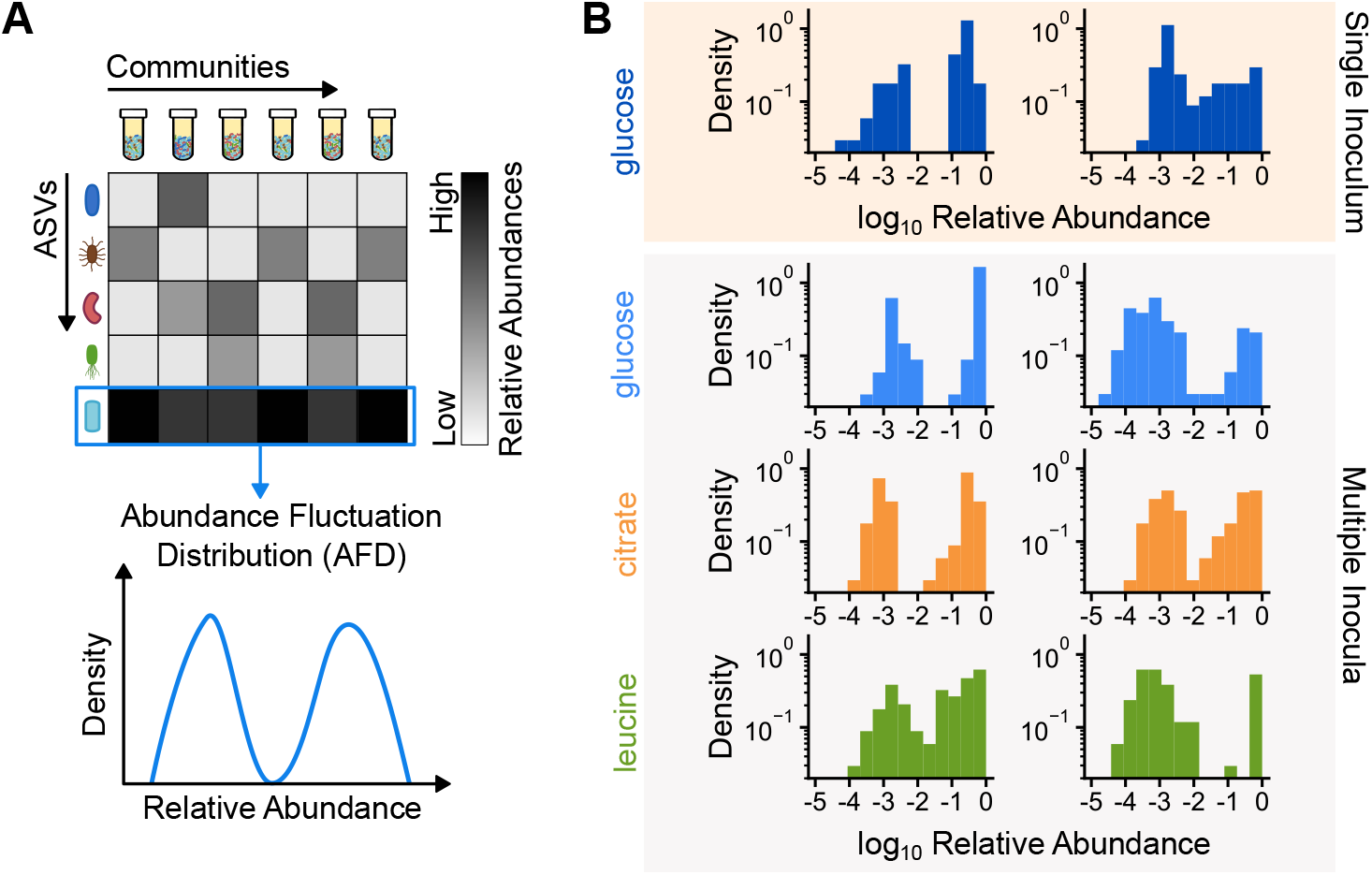
Most ASVs have bimodal Abundance Fluctuation Distributions (AFDs). (A) Schematic illustrating the construction of an AFD for a single ASV from its relative abundances across replicate communities. (B) Empirical AFDs for representative ASVs from single and multiple inocula experiments, showing consistently bimodal patterns.

In this work, we considered replicate *in vitro* communities assembled from one or multiple inocula using the same medium under a serial dilution protocol [11, 12]. Here, we consider the communities assembled after 18 dilution cycles for the single inoculum and 12 dilution cycles for the multiple inocula. Figure 1B shows that ASVs exhibit a bimodal AFD in these experimental communities. A bimodal AFD implies that there are two typical abundances, corresponding to the two modes of the AFD. An ASV is present in high abundance in some samples and low abundance in other samples.

In Supplementary Information **??**, we consider several metrics to evaluate and quantify the degree of bimodality. We focused on the log of the likelihood ratio between a unimodal and a bimodal model (see Methods), which quantifies the information lost by approximating the empirical distribution with a unimodal vs. a bimodal model. In order to evaluate the degree of bimodality in the data, we compared the empirical value of difference of the log likelihood ratio Δ ℒ_*d*_ between the empirical and the one obtained from a null model (see Methods). Our analysis revealed that ASVs in communities assembled from a single inoculum exhibit log-likelihood ratios larger than what would be expected by chance under a unimodal distribution (Figure **??**), demonstrating that the vast majority of ASVs display a bimodal AFD. This finding extends previous observations [12] that characterized bimodality for the two most abundant genera (*Alcaligenes* and *Pseudomonas*).

### The effects of compositionality are not the source of the observed bimodality

Observed bimodality can be due to compositional effects. Since AFDs deal with relative abundances rather than absolute counts, compositional effects could potentially induce bimodal patterns, as a positive (negative) fluctuation of absolute abundance of a single ASV could induce negative (positive) fluctuations of the relative abundances of all the other ASVs, making them display an apparent bimodality in relative abundance. To address compositional effects as a potential source of bimodality, we generated *in silico* data with variable proportions of truly bimodal ASVs to test the effect of compositionality in inducing apparent bimodality in relative abundances when true absolute abundances are not bimodal (see Supplementary Information **??**).

We observe that compositional effects can indeed influence the shape of the relative abundance distributions. The key parameter determining the presence of these effects is the the coefficient of variation of absolute abundance: when bimodal ASVs have much higher coefficient of variation than unimodal ASVs, compositional effects are more likely to create artificial bimodal patterns.

Since direct measurements of absolute abundances are rarely obtained in sequencing-based studies, we first explored which common methods of removing compositional effects could reliably recover the true form of the distribution generated by our toy model. We found that additive log-ratio (alr) transformation using unimodal ASVs as references successfully revealed the true underlying bimodal distributions, as bimodal ASVs remained bimodal and unimodal ASVs returned to their true unimodal shape. In contrast, other methods such as the centered log-ratio (clr) that uses the geometrical mean as a reference, fail in removing those effects due to the geometrical mean itself being bimodal, clustering the transformed data around the peak of the geometrical mean. While implementing the alr-transformation is straightforward, we were faced by the challenge of determining the reference ASV. Given the absence of prior prior information on the unimodality/bimodality of the true abundance, we investigated whether the abundance of a single ASV can induce apparent bimodality in remaining ASVs. Figures **??**-**??** show that there is no case that a single ASV can induce observed bimodality in the alr-transformed data.

In addition to the test above, for the single inoculum dataset, we also estimated absolute abundances by multiplying relative abundances by optical density measurements. These estimated absolute abundances still displayed bimodality (Figure **??**), further confirming that the observed bimodality in relative abundances is not just a consequence of compositional artifacts.

### The abundance of each ASV can be assigned to two distinct states

A key implication of bimodal AFDs is that each ASV exists in one of two distinct states: low- or high-abundance. Assigning an ASV in each assembled community to one of these two states involves a certain degree of uncertainty. We considered a mixture model to define the probability that the abundance of a species in a particular sample belongs to either of the two states (Figure 2A and Section).

**Figure 2:**
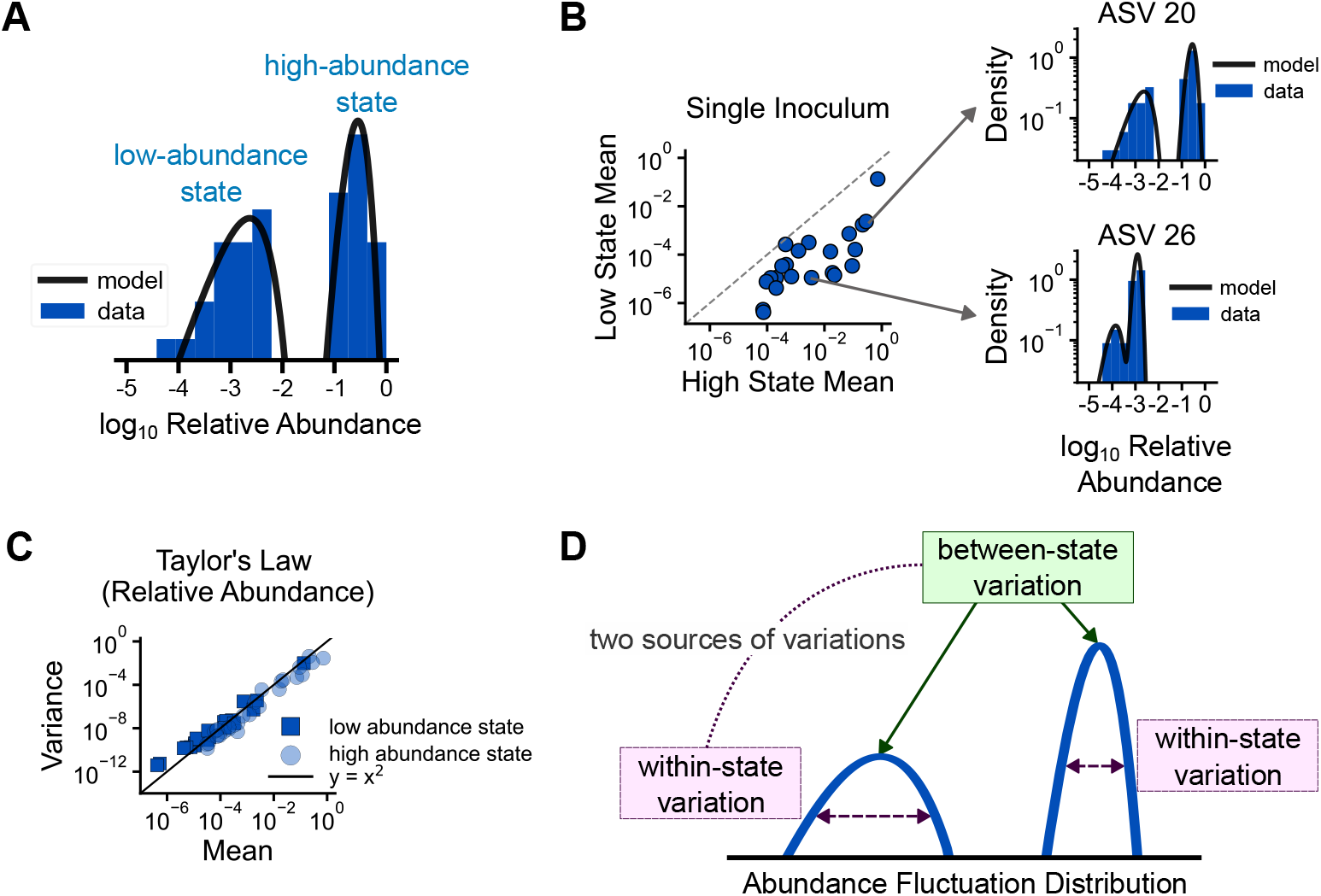
The total variation of the AFD of a given ASV is determined by between-state variation and within-state variation. (A) A bimodal AFD implies the presence of two states: low- and high-abundance. Here we show the AFD of one of the ASVs from the single inoculum experiment modeled using a mixture of two gamma distribution. (B) Comparison of mean relative abundances between low- and high-abundance states for all ASVs in the single inoculum dataset (left). Example comparison of two ASVs showing that low-abundance states do not correspond to the rarest ASVs (right). (C) Taylor’s Law with exponent 2 holds for each state across all ASVs in the single-inoculum dataset; the same pattern is observed in the remaining three datasets (Figure **??**). (D) Schematic illustration of a mixture model that fits AFDs (blue line). There are two sources of variation in the AFD: between-state variation and within-state variation.

Each taxon is therefore characterized by two typical abundances, corresponding to the low- and high-abundance states. Figure 2B and Figure **??** show that the values of these low- and high-states are unique to each ASV. For instance, in the communities assembled from a single inoculum, the typical relative abundance in the low-abundance state and high-abundance state for ASV 20 is 0.002 and 0.28 while the typical relative abundance in those states for ASV 26 is 0.0001 and 0.001. These examples not only show the uniqueness of the two states, but also imply that an ASV in a low-abundance state is not necessarily “rare” in absolute terms, as its relative abundance can exceed the high-abundance state of other ASVs. Therefore, low-abundance states cannot be dismissed as being excessively close to the lower limit of observation.

One could ask whether low-abundance states correspond to an experimental or technical artifact, due to transient or non-growing populations. The serial dilution design of the original experiment provides a framework for distinguishing true low-abundance states from contaminants or transients, which would disappear after a low number of transfer cycles. Specifically, random sampling occurs at each step of the dilution procedure, meaning that only ASVs that are present at sufficient abundances will persist through multiple transfer cycles (Figure **??**). The consistent observation of low-abundance states after a transfer has reached steady-state suggests that community members have multiple attractors rather than being transient members in a fraction of the replicate communities.

### The bimodal abundance fluctuation distribution is well described by a mixture of two gamma

Previous work in natural [2, 23, 24] and experimental [4] communities showed that Abundance Fluctuation Distributions (AFD) are well described by a gamma distribution. This distribution can be interpreted as the stationary distribution of an underlying model of limited growth in a fluctuating environment (the Stochastic Logistic Model [1, 2]).

Building on these insights, we modeled the empirical ASV bimodal abundance distributions using a mixture of two gamma distributions (Section ). We tested the performance of the two-gamma mixture model against the unimodal gamma. The mixture of two gamma distributions more accurately reproduces the empirical AFDs compared to the unimodal gamma model (see Supplementary Information **??**).

The AFD of each ASV is therefore characterized by five parameters: the probability that the ASV is in low-abundance state and the mean and coefficient of variation of abundance in each of the two states. Figure **??** - Figure **??** show that the mean abundance of an ASV in the high-/low-abundance state is uncorrelated with the probability of being in that state. As a generalization of what was observed for unimodal AFDs [2, 24], Taylor’s Law holds independently for both of the two states, as the coefficient of variation is independent of the average abundance for both the high- and the low-abundance state (Figure 2 and Figure **??**).

The mutual independence of the mean abundance and the probability of being in the high-/low-abundance state suggests that there are two (macro)ecological sources of variation shaping fluctuations of abundance across communities. One source of variation determines whether an ASV is in one of the two states, which we refer to as between-state variation. The other source of variation determines the fluctuations of abundance within each state, which we refer to as within-state variation (Figure 2D).

### Between-state variation drives abundance correlations across ASVs

As a result of the effect of several ecological processes, we expect ASVs to have correlated fluctuations. For instance, limiting similarity (e.g., through competition) determines negative correlations between ASVs, while environmental filtering produce positive fluctuations. Both types of variation (within- and between-state) determine the abundance of ASVs in a given community. In order to disentangle their ecological significance, we explore how they independently contribute to the overall correlations of ASVs.

Figure 3 compares the empirical pairwise abundance correlations across ASVs with two null models that aim to disentangle the influence of within- and between-state variation. The abundance correlation across ASVs concentrates around zero with small peaks at both tails, implying strong correlations for some pairs of ASVs. For the first model, we shuffled the abundance of each ASV within its state (i.e., low-abundance state or high-abundance state), retaining between-state information. This null model successfully reproduces the observed distribution of correlations. Meanwhile, for the second null model, we shuffled abundances across communities, removing both within- and between-state variation. This null model fails to reproduce the empirical correlation distribution. The correlations are therefore mostly due to between-state variation (whether species co-occur in the same low-/high-abundance state in a given sample or not) and not to the actual abundances they have in each state.

**Figure 3:**
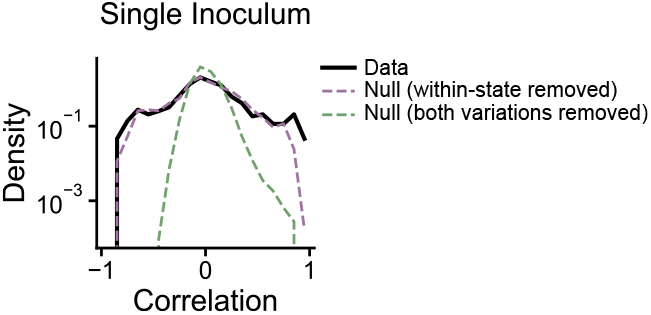
Between-state variation drives abundance correlations across ASVs. The empirical distribution of pairwise ASV abundance correlations for the single inoculum dataset is compared to two null models. A null model that removes within-state variation reproduces the empirical correlations while a null model that removes both within- and between-state variation fails to reproduce the empirical correlations. Similar patterns are also observed for multiple inocula datasets (Figure **??**). This demonstrates that correlations are driven by whether ASVs are jointly in their high- or low-abundance states.

To further strengthen this result, we explored the abundance correlations for pairs of ASVs conditioned on their states (e.g., the correlation between ASV 1 and ASV 2 conditioned to ASV 1 being in high-abundance state and ASV 2 in low-abundance state). We generated an *in silico* dataset by drawing abundances of each ASVs from the mixture of two gamma distribution with the empirical fitted parameters and sampling discrete sequence counts using the empirical total number of reads. These generated data sufficiently capture empirical correlation distributions (see Figure **??**). We conclude that within-state variation does not contribute substantially to the correlation structure observed across ASVs.

### Between-state variation is strongly correlated across ASVs

Since the within-state variation appears to be independent across ASVs, we explored the degree of correlation of the between-state variation. We introduced a new variable *σ*_*i*_, defined to be equal to ±1 if ASV *i* is found in the high- or low-abundance state, respectively (Figure 4A and Method). We then calculated the correlations of *σ*^*i*^ for each pair of ASVs across communities and compared the empirical correlation distribution with a null model obtained by shuffling the variable 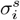 for each ASV across communities. This null model preserves fraction of communities where an ASV is found in a given state, while removing their associations. The null distribution differs significantly from the empirical distribution (Figure **??**), since data have a more pronounced tail for both positive and negative correlations. This observation implies that the between-state variation strongly drives the correlation between ASVs.

**Figure 4:**
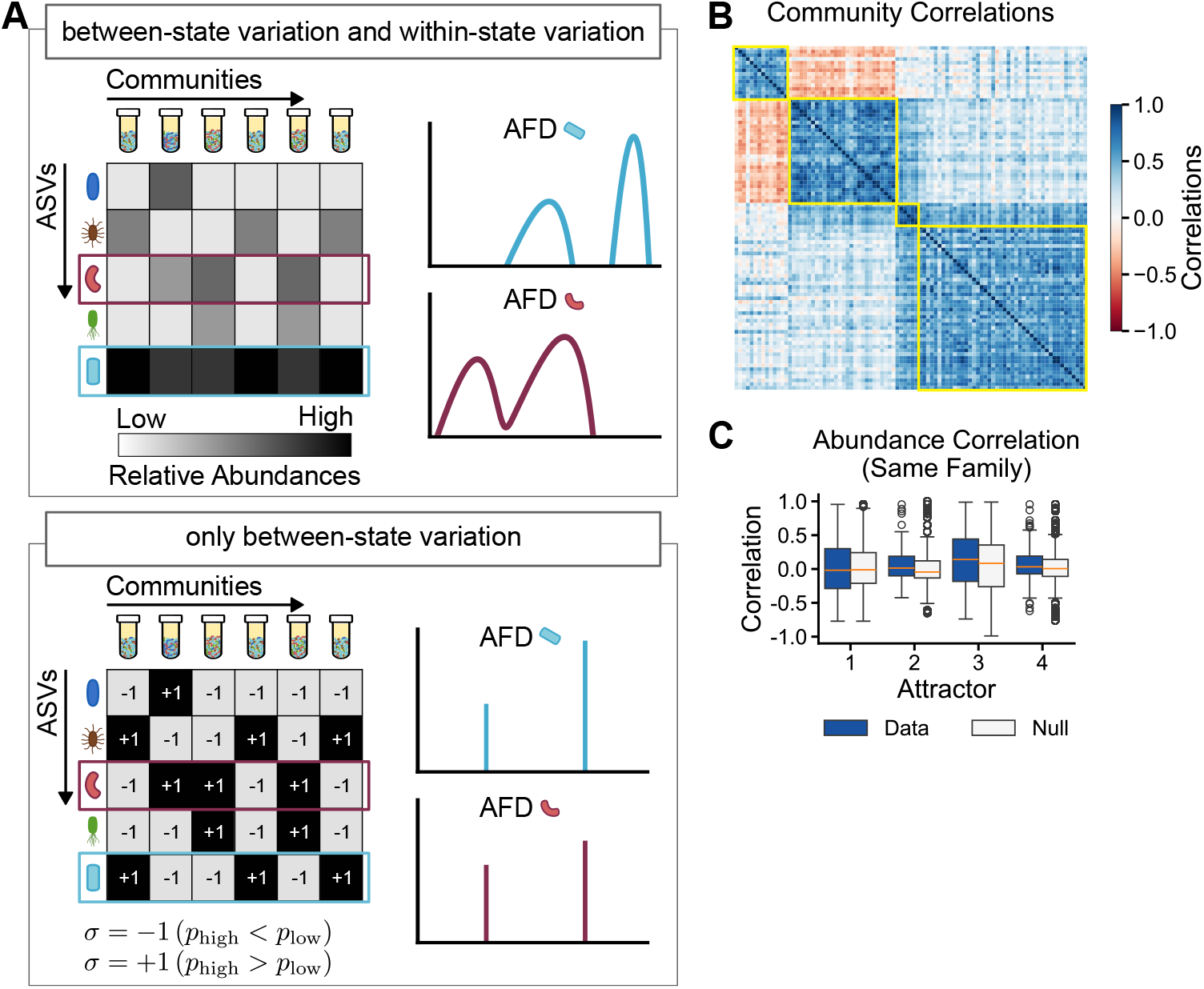
Between-state variation correspond to alternative stable states. (A) Schematic showing the conversion of continuous relative abundances into a binary state variable (*σ* = −1 for low, +1 for high) that captures only between-state variation. (B) The community correlation matrix based on the binary state variable reveals a distinct block structure, identifying four alternative stable states (attractors). (C) Within each attractor, abundance correlations for ASV pairs from the same family do not differ from a null model, indicating no taxonomic signal at this scale.

### The correlated variation between states reflects the presence of multiple attractors

Building on prior findings on alternative taxonomic compositions as well as functional states in experimental microbial communities grown on a single carbon source [12], we investigated whether the correlated between-state variation in the ASV-level reflected the presence of alternative stable states. We calculated the similarity between communities by using the binary assignment (*σ*_*i*_), capturing the fraction of ASVs appearing in the same state in two communities.

By reordering the communities using hierarchical clustering, a block structure clearly emerges in the correlation matrix (Figure 4B). This block structure is robust to the choice of clustering method and threshold (Figure **??**). Pairs of communities within the same block (group) show high positive correlation, indicating that they typically co-occur in the same high-/low-abundance state across communities. Furthermore, communities within the same group show consistent correlation patterns when compared with communities from different groups. This block structure likely reflects the existence of alternative stable states corresponding to different groups of communities. In particular, for the single inoculum case, we reveal a non-trivial correspondence with previously documented taxonomic and functional compositions [12] (Figure **??**). Although these identified attractors do not align perfectly with the compositional classification, they capture consistent groupings of communities with similar composition.

We further validated these structural patterns using additional similarity metrics. The binary correlation matrix shows strong alignment with alternative measures of similarity (complement of Bray-Curtis and Jaccard indices, see Figure **??** and Figure **??**).

The block structure is observed both in the case of single inoculum and multiple inocula (Figure **??**). Therefore, the structure cannot be uniquely explained by an inoculum effect. However, in the case of multiple inocula, communities assembled from the same inoculum are more likely to cluster in the same attractor (Figure **??**). This effect is driven by the fact than not all ASVs are detected in all inocula. Several ASVs appear only in communities assembled from specific inocula and are consistently not detected in others (Figure **??**). This result shows that one can not simply treat different inocula as exchangeable. Because of this effect, for the following analysis we focused on the single inoculum case.

### ASVs abundance is not constrained by family abundance within attractors

What is the most relevant phylogenetic or taxonomic scale at which to define ecological populations is a fundamental issue in microbial ecology [25]. It has been previously proposed that the family level is the finest taxonomic scale where compositional variation becomes reproducible in experimental communities assembled under fixed conditions [11]. Under this view, composition at the ASV level lacks reproducibility even in controlled conditions while family-level abundance is reproducible.

In the data we consider how the existence of multiple attractor slightly alters this picture of reproducibility in community assembly. We expect family abundance to be reproducible within each attractor, while it could vary across attractors as consequence of the variable state of a community [12]. The reproducibility of the abundances of families, together with the variability at the ASV level, would imply ASVs within the same family typically have negative correlations.

Figure 4C shows that the values of the abundance correlations of pairs of ASVs within the same family do not differ than what is expected by a null model (obtained by shuffling taxonomic assignments while preserving abundance compositions). This lack of evidence of negative correlations is consistently observed across attractors. We also observe the same pattern when, instead of abundance, we considered the correlation of state occupancy (*σ*) of ASVs within families (Figure **??**. The fluctuations of abundance within an attractor do not display a phylogenetic or taxonomic signal.

### ASVs groups with same and reciprocal prevalence

To characterize the multiple attractors in the communities, we calculated the probability that a species is in its high-abundance state given a particular attractor, which we refer to as prevalence. Prevalence provides a meaningful characterization of the block structure by identifying which ASVs consistently occupy high or low-abundance states within each attractor.

ASVs exhibit different prevalence values across the four observed attractors. Prevalence was computed as the proportion of samples in a given attractor in which an ASV was in the high-abundance state (see Methods for details). For example, an ASV that is predominantly in the high-abundance state across samples within an attractor will have high prevalence, whereas an ASV that is predominantly in the low-abundance state will have low prevalence. Each ASV has a characteristic prevalence profile, which can vary substantially across attractors. Some ASVs have similar prevalence values across attractors, while others show distinct variation. Figure 5A illustrates these prevalence patterns across all ASVs (see Figure **??** for detailed ASV identities).

**Figure 5:**
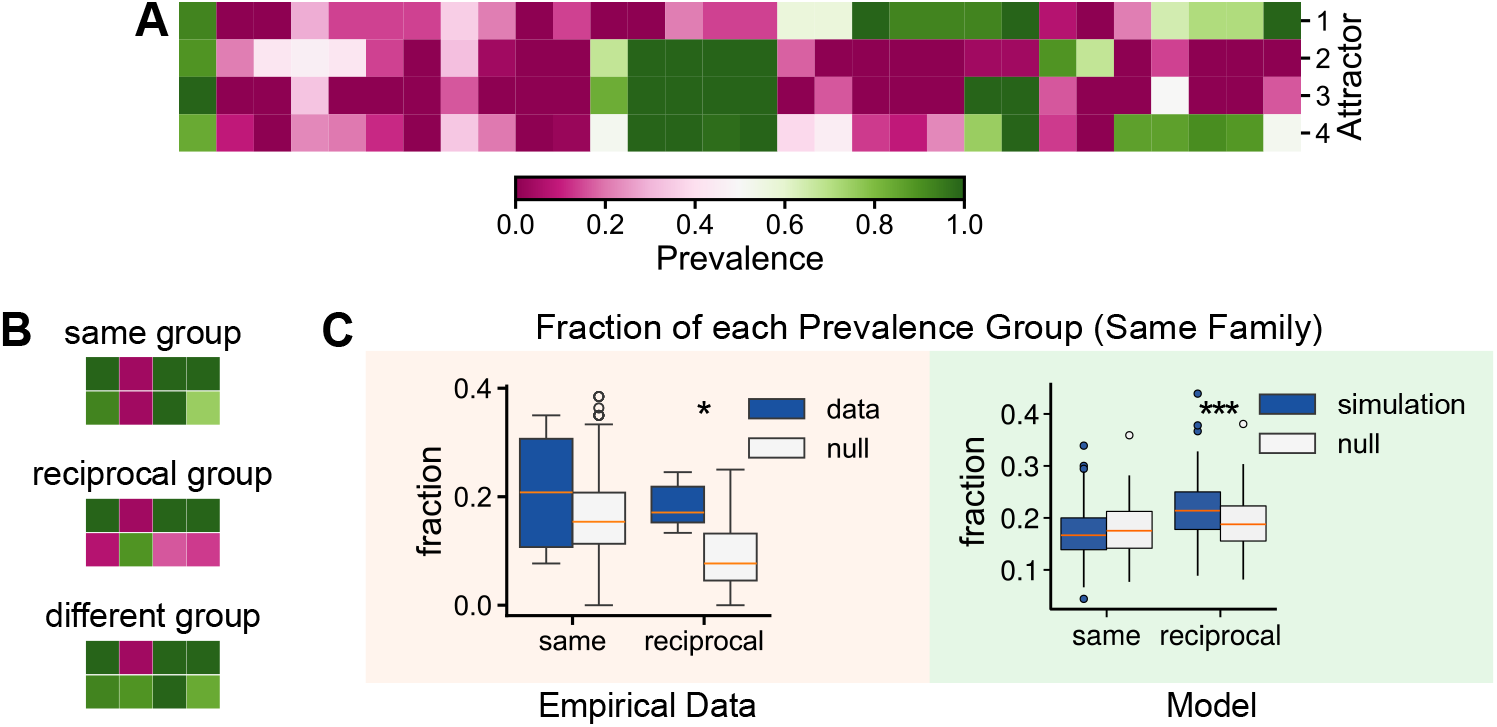
Phylogenetically close ASVs exhibit reciprocal prevalence patterns across attractors. (A) Heatmap showing ASV prevalence across the four attractors. Each column represents an ASV; each row represents an attractor. Colors indicate prevalence (the proportion of samples in which an ASV is in the high-abundance state). ASV identities are provided in Figure **??**. (B) Examples of ASV pairs belonging to the three categories: same (similar prevalence), reciprocal (inverse prevalence), and different (neither similar nor inverse) groups. Each row represents an ASV and each column represents an attractor; the layout is rotated relative to panel A for clarity. (C) *Left* : Compared to a null model, phylogenetically close ASV pairs (within the same family) are significantly more likely to have reciprocal prevalence patterns (p-value *<* 0.01 (KS-test)), suggesting state-dependent competitive exclusion differentiates the attractors. *Right* : Simulations of the Ising–Hopfield model for medium environmental filtering (*q* = *M*) and high competitive exclusion (*p* = *H*). This parameter combination is the only one that reproduces the empirical pattern of statistical significance of fraction of ASVs in the reciprocal group (see Figure **??.**

Despite the theoretical possibility of 16 prevalence patterns (2^4^ combinations) across four attractors, only 8 distinct patterns were observed, with 80% of ASVs belonging to just four of these patterns. This indicates a simpler underlying structure than if each ASV responded independently. Pairwise comparisons of ASV prevalence profiles reveal interesting relationships that fall into three distinct categories (Figure 5B). ASV pairs are classified as same if they exhibit similar prevalence patterns across attractors (e.g., both having high prevalence in attractors 1, 3, and 4, but low prevalence in attractor 2). Pairs are classified as reciprocal if they show inverse patterns: one ASV has high prevalence where the other has low, and vice versa across attractors. When multiple ASVs share a prevalence pattern and multiple ASVs share the opposite pattern, any pair formed between these two groups is considered reciprocal. Pairs that fit neither the same nor reciprocal category are classified as different.

### ASVs within the same family are more likely to have reciprocal prevalences

We found that pairs of ASVs within a family do not have correlations of abundance or state different than what is expected by chance. On the other hand, we showed the existence of clear prevalence patterns shared by multiple ASVs. We therefore asked whether prevalence patterns have a detectable taxonomic signal. More specifically, we asked whether ASVs within the same family are more/less likely than what expected by chance to have the same or the reciprocal pattern.

We can assume that different attractors correspond to different “realized” environmental conditions which favor different sets of community members. A scenario dominated by environmental filtering would imply similar species (ASVs within the same family in this case) to be more likely to have the same prevalence, as they would both thrive in the same conditions. On the contrary, an attractor could favor one ASV over another, thereby determining which ASV is favored among closely related ASVs.

Figure 5C (left) shows the fraction of pairs of ASVs that have the same or reciprocal prevalence patterns. This number is around 20% across attractors for both same and reciprocal pairs. To investigate whether these fractions are large or small, we compared them with a null model obtained by shuffling the taxonomic assignment of ASVs while preserving the prevalence matrix of Fig. 4A.

The null model reproduces the observed fraction of pairs that have the same prevalence pattern (KS-test, p-value=0.47). On the other hand, the null model predicts a fraction of pairs with the reciprocal prevalences significantly lower than what was observed in the data (KS-test, p-value*<* 0.01). Taxonomically similar ASVs (i.e., within the same family) are ~ 2 times more likely to have reciprocal prevalences than what expected by chance.

It is interesting to notice that, as shown in Figure. 5C (left), a negative correlation between ASVs states is not observed: in any particular attractor two species within the same family are not in a different abundance state more often than expected by chance. On the other hand, when comparing different attractors we observe that the prevalences are more likely to be negatively aligned than expected by chance.

### A minimal model explains the structure of alternative stable states

To understand how the patterns of community structures we observed could arise from basic ecological rules, we developed a minimal mathematical model, by adapting a framework originally from statistical physics (the Ising–Hopfield model [26]). Our model incorporates two fundamental ecological drivers: (i) environmental filtering, which pulls the community towards specific compositional states, and (ii) competitive exclusion, which prevents ecologically similar species from dominating simultaneously.

Consistent with the observation that species abundances are bimodal, we model each ASV not as a continuous variable, but as a simple binary switch between a low- and a high-abundance state. The goal of the model is to determine which combinations of these ecological forces are capable of reproducing our empirical observations. In particular, under what conditions do we expect similar species (e.g., species of the same family) to have reciprocal prevalences, as observed in Figure 5C (left).

In order to model our observations it is necessary to mathematically define what we mean by “attractor”. We define attractors to be the configuration of a community with the lowest incompatibility among ASVs with different preferences, determining the stability of the community. This definition is analogous to that of energy landscapes in physics: community dynamics drive the systems toward configurations with the lowest energy, corresponding to stable states (“attractors”).

We define the probability of observing a community configuration (a given sequence of low/high-abundance states) as the result of the tension between two forces (see, Appendix **??**, and Fig. **??**). The first force represents environmental filtering. We assume the environment corresponds to specific “templates” compatible species combinations, ultimately based on their traits. The community composition is statistically more likely to align with one of these templates. This mechanism creates the alternative stable states we observed experimentally: distinct, reproducible configurations that the community settles into. The second force accounts for competition between closely related species. We assume that two species of the same family do not tend to be in a high abundance state at the same time.

These two forces, therefore, create opposing ecological tension. The first force tends to select similar species to be in the same abundant state, as they share similar traits. Competitive exclusion, on the other hand, disfavors such configurations. We quantify these two effects as two parameters (*p* and *q*, both in the interval [0, 1]). The value *q* captures the level of similarity of the environmental templates for ASVs belonging to the same family, relative to two AVSs chosen at random. The parameter *p* quantifies how strong the exclusion/competition between ASVs of the same family is, relative to what is observed on average. The model reproduces the existence of multiple attractors, with distinct prevalence patterns as shown in Figure 5A (see Figure **??**A). By analyzing simulations covering the entire parameter range of *p* and *q*, we find that only a combination of moderate environmental filtering and high competitive exclusion is consistent with the data (Figure 5C (right) and Figure **??**B-C). This result indicates that both filtering and exclusion play a non-negligible role in shaping community composition, with competitive exclusion at the ASV-level due to limiting similarity being the dominant force.

## Discussion

In this study, we dissected the fine-grained structure of experimentally assembled microbial communities and uncovered a general pattern characterizing their variability: the bimodal abundance distribution of individual ASVs. This finding stands in sharp contrast to the unimodal distributions typically observed in diverse natural ecosystems [2, 4] and provides a new framework for understanding the emergence of alternative stable states in microbial consortia. Our results demonstrate that community structure is not primarily driven by continuous fluctuations in abundance, but by discrete, binary-like switching of constituent taxa between high- and low-abundance states. Such results are reminiscent of what is observed in long timeseries of the human gut microbiome [19], where individual microbial populations (e.g., OTUs) are observed to abruptly change in their typical abundance, suggesting the existence of underlying alternative stable state.

The most critical finding of our work is that nearly all ASVs exist in one of two distinct modes. This bimodality is not a compositional artifact but an intrinsic property of these communities, suggesting the presence of strong nonlinear dynamics and feedback loops. In the controlled environment of a serial dilution experiment, such bimodality likely arises from ecological interactions like cross-feeding or priority effects, which create two distinct basins of attraction for each population. An ASV may either successfully establish and reach a high carrying capacity, or fail to establish and persist at a low, suppressed level. The abundance of an ASV is therefore better described not by a single mean value, but by the properties of these two states and the probability of occupying them. It is worth noting that the probability of multiple stable states emerging may ultimately be something that can be manipulated. For example, alternative stable states in the experimental communities examined here and elsewhere were provided with a single supplied substitutable resource [11, 12, 27], potentially reflecting the role of resource composition. In addition, our communities were assembled from an initial soil sample harboring extremely high diversity (~ 10^3^ ASVs) and it has been recently found that alternative stable states tend to emerge with increasing initial richness in complex environments [28]. These ecological features provide avenues of exploration for manipulating alternative stable states in experimental and natural communities.

By decomposing the total variation into “within-state” and “between-state” components, we showed that the ecologically significant information is captured by the latter. The correlations between ASV abundances across communities were almost entirely explained by which state (high or low) the ASVs were in, not by their precise abundance fluctuations within those states. This discovery allowed us to develop a novel, bottom-up method for identifying community-level attractors based on the collective binary states of all ASVs. This approach revealed a clear block-like structure of alternative states, providing a more granular view than previous analyses based on overall community similarity or family-level composition [12].

Our analysis provides a compelling resolution to the family-level reproducibility and ASV-level variability. While previous work [11] highlighted functional redundancy within families as a key organizing principle, our results reveal a more competitive and structured outcomes of ecological dynamics. We found that ASVs within the same family are significantly more likely to exhibit reciprocal prevalence patterns across attractors than expected by chance. That is, if one ASV thrives in a particular attractor, its close relative is likely suppressed, and vice-versa in a different attractor. Our result strongly suggests that different attractors represent distinct ecologies that favor one close competitor over another. Rather than being functionally interchangeable, these closely related ASVs appear to be engaged in state-dependent competition, where the identity of the “winner” is contingent on the collective state of the entire community.

In conclusion, our work reframes the analysis of microbial community structure by shifting the focus from continuous abundances to discrete, bimodal population states. This perspective not only provides a powerful, data-grounded method for identifying alternative stable states but also reveals that fine-scale niche differentiation among close relatives is a fundamental mechanism driving community assembly. Future work should aim to uncover the specific interactions that give rise to these bimodal distributions and reciprocal patterns, and to explore whether this binary-state framework can be extended to understand the more complex dynamics of natural microbial ecosystems.

## Methods

### Data

We analyzed experimental data of the assembled community of soil bacteria in a single-limiting carbon source with a single inoculum [12] and multiple inocula [11]. For the single inoculum data, we refer to the data used in [29] for the 18 transfers without migration. We reprocessed the multiple inocula data to include singletons using DADA2 pipeline (version 1.30.0) to infer Amplicon Sequence Variants (ASVs) [30]. The forward and reverse reads were trimmed to 220 and 160 nucleotides, respectively and the reads below 160 and above 251 nucleotides were discarded. We used “consensus” method in DADA2 to remove chimera. SILVA version 138 was used to assigned taxonomy to ASVs.

### Abundance Fluctuation Distribution Models

Our observation of bimodal patterns in ASVs’ AFDs motivated this exploration of a mixture model, which extends beyond the unimodal distribution framework used in previous study [2]. Here, we first re-introduced the unimodal model and the extension into the mixture model.

In the previous study, AFDs are gamma distributed. For ASV *i*, the distribution of the relative abundance *x*_*i*_ is

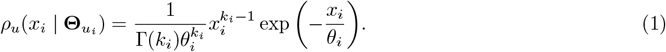

There are two parameters in the distribution above 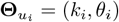 with *k*_*i*_ is the shape parameter and *θ*_*i*_ is the scale parameter. Previous study [2] used mean relative abundance of ASV 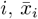, and inverse of the square of coefficient of variation *β*_*i*_ to parameterize the distribution. We could easily convert the two notation of parameters by the relations 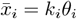, and, 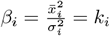 since 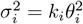.

The gamma distribution provides a continuous model for relative abundances. However, the sequencing data consists of discrete read counts that are subject to sampling effects. Under the assumption that the relative abundances follow a Gamma distribution and reads are sampled randomly, the resulting read counts follow a negative binomial distribution [2]. Specifically, the probability of observing *n*_*i*_ reads for ASV *i* in a sample *s* with *N* total number of reads is

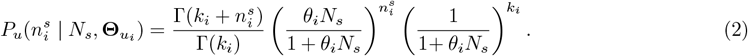

Since our observation shows bimodal AFDs, we extend a single gamma distribution into a mixture of two gamma. Therefore, the distribution of the relative abundance of ASV *i, x*_*i*_ is

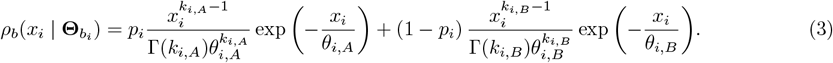

with *p*_*i*_ being the mixing parameter of the distribution and 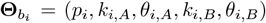. The parameters *k*_*i,A*_, *θ*_*i,A*_ and *k*_*i,B*_, *θ*_*i,B*_ can be linked to the mean and variance of the relative abundance of each of two modes. Here, we assigned *A* as the mode with lower typical abundances and *B* as the mode with higher typical abundance. Similarly, the probability of observing a read count in a sample, 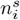 is the convolution of this mixture and sampling distribution which results in a mixture of two negative binomials

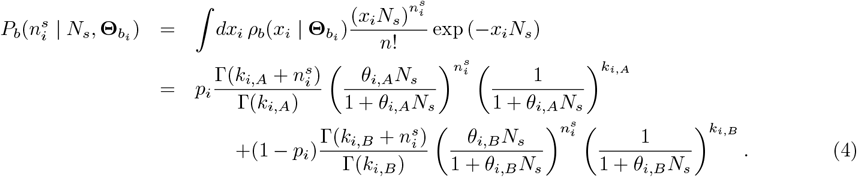

### Parameter Estimation with Maximum Likelihood Estimation

We estimated the parameters of the gamma distribution and mixture of two gamma distribution for each ASV independently. We used a Maximum Likelihood Estimator (MLE) in which we maximized the log-likelihood of the read counts. Estimation of the parameters of the gamma distribution was done not to fit the AFDs rather to obtain parameters that were used to generate null model for our statistical analysis.

For the unimodal distribution, the log-likelihood of the read counts is

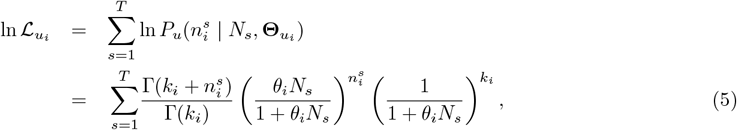

and for the bimodal distribution, the log-likelihood is

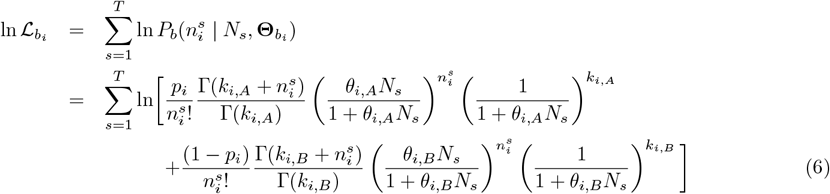

where *T* is the total samples. We could rewrite Equation 6 as

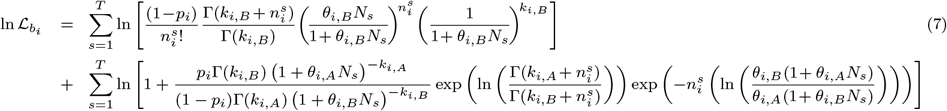

We solved the optimization of the log-likelihood numerically by minimizing the negative of the log-likelihood using SciPy package in Python 3.11.4 with Nelder-Mead algorithm. Initial value of the parameters of each ASV for the unimodal distribution were obtained using empirical mean *x*_*i*_ and variance *σ*_*i*_. Both value then converted into *k*_*i*_ and *θ*_*i*_ as shown in the previous section. For the bimodal distribution, the estimation of the two modes of the distribution of each ASV was done using k-means clustering. The clustering separated the read counts into two clusters. Initial value of *p*_*i*_ were estimated from the proportion of samples that belonged to the mode with lower mean after clustering. The initial value of the parameter of each mode: *k*_*i,A*_ and *θ*_*i,A*_, *k*_*i,B*_ and *θ*_*i,B*_, was estimated from the empirical mean and variance of each mode after clustering.

One issue with the optimization is that the algorithm can become stuck at local minima. To overcome this, we used various initial values and choose the one with the smallest negative log-likelihood as our estimator. We varied the initial conditions by doubling or halving the variance of the initial condition above while keeping the mean. For the bimodal distribution, the doubling or halving was done for each mode separately and for both modes simultaneously. We also used the parameters obtained after minimization with the previously described initial values as initial condition, as well as doubling and halving the variance for these parameters. For any ASV where the optimization did not converge, we estimated that parameter as follows: we used the empirical mean and the average of the coefficients of variation of relative abundances for ASVs where estimation was successful (converged) to obtained the shape and scale parameters.

### Generating Datasets from Fitted Parameters

To generate a dataset with unimodal distribution, we kept the same number of ASVs and replicates as in the empirical data. For each replicate, we drew the relative abundances of each ASV independently from Equation 1 using the best-fitted parameter from the Maximum Likelihood Estimation of equation 5 (Section ) and constrained the relative abundances to sum to one. We then performed multinomial sampling with the drawn relative abundances and the total number of read counts from the corresponding sample in the original data.

For the generated bimodal datasets, we followed the same procedure but drew relative abundances from the two-gamma mixture distribution (Equation 3) using parameters fitted from the bimodal model (Equation 7).

### Likelihood Ratio Test

For each ASV *i*, the log of the likelihood ratio (ΔL_*i*_) is defined as

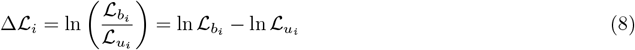

where 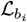 is the likelihood of the bimodal model (Eq. 6) with best fitting parameters and 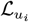 is the likelihood of the unimodal model (Eq. 5) with best fitting parameters.

We constructed the null model by generating 300 datasets (*S* = 300) with unimodal distribution (see Section ). We fitted each of ASV *i* in each of the generated dataset *s* with bimodal model (Equation 7) and unimodal model (Equation 5), similar to the empirical data. Then, for each ASV, we computed log of likelihood ratio of the ASV in the generated dataset Δℒ_*i,s*_ and taking the average over all generated datasets as follows

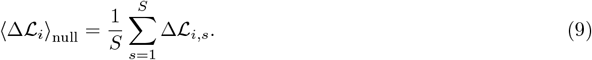

### Assignment of Low Abundance States and High Abundance States

Our two-gamma model enables us to define two states for ASVs in a sample: high- and low-abundance. An ASV is assigned to the low-abundance state if its relative abundance, *x*_*i*_, belongs to the first term in Equation 3, while an ASV in the high-abundance state has a relative abundance belonging to the second term (Methods. However, state assignment is non-trivial when the two distributions overlap, as abundances in the overlap region could potentially be assigned to either distribution. To address this, we introduce a confidence measure that quantifies how certain we are that a given relative abundance belongs to one of the gamma distributions

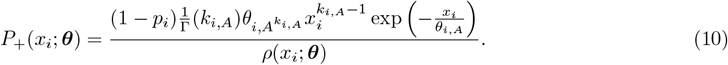

This quantity represent the probability that relative abundance *x*_*i*_ belongs to the mode with higher typical abundances. Thus we also have *P*_−_(*x*_*i*_; ***θ***) = 1 − *P*_+_(*x*_*i*_; ***θ***) that represent the probability that the relative abundance *x*_*i*_ belongs to the mode with higher relative abundance. We then constructed a function that maps the relative abundance of ASVs to the states as

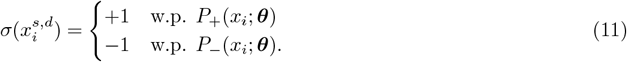

Assigning a state of an ASV with relative abundance *x*_*i*_ in a sample *s* can be seen as one realization of drawing the variable 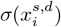. If the procedure is repeated, then different values will be obtained. Because of that, we took the average of 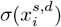 over the realizations

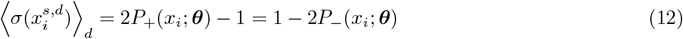

The assignment of the relative abundance of ASV *i* in sample *s* to the low- or high-abundance states is then determined by 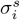

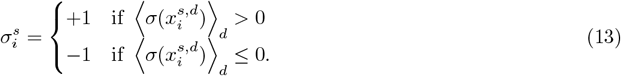

### Community state clustering and ASV prevalence

Communities were clustered based on their pairwise correlations, calculated as the fraction of ASVs in the same abundance state between any two samples. Hierarchical clustering was performed using complete linkage with a distance threshold of 0.5 times the maximum dendrogram distance, identifying four distinct community states (attractors). Alternative clustering methods (Ward linkage, different thresholds) were used to assess the robustness of the block structure. Correlation and clustering analyses were performed using standard implementations in SciPy.

For each ASV *i* in attractor *α*, prevalence was defined as the fraction of samples within that attractor in which the ASV occupied the high-abundance state:

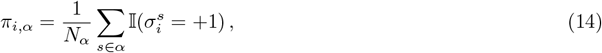

where *N*_*α*_ is the number of samples assigned to attractor *α*, and I(·) denotes the indicator function. The prevalence, *π*_*i,α*_, therefore quantifies the probability that ASV *i* is in the high-abundance state within attractor *α*.

To compare prevalence patterns across ASVs, prevalence values were discretized into high- and low-prevalence categories. An ASV was classified as having high prevalence in attractor *α* if *π*_*i,α*_ ≥ 0.5, and low prevalence otherwise. Pairwise relationships between ASVs were then classified based on their binary prevalence profiles across attractors. Two ASVs were assigned to the same category if they shared identical prevalence classifications in all attractors. Two ASVs were assigned to the reciprocal category if their prevalence classifications were opposite in all attractors, such that one ASV was high where the other was low. ASV pairs that satisfied neither condition were classified as different.

### Theoretical framework

To mechanistically interpret the observed alternative states, we employed a minimal Ising-Hopfield model where each ASV adopts a binary abundance state *σ*_*i*_ ∈ {−1, +1}. The system’s energy landscape is shaped by environmental filtering, modeled as Hopfield’s “memory patterns” *ξ*^*µ*^ that define basins of attraction. We parameterized the taxonomic structure of these environments using a filtering parameter *q* ∈ [0, 1]. This parameter governs the probability that ASVs within the same family respond coherently to an environmental axis 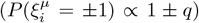; thus, *q* tunes the system from random environmental selection (*q* = 0) to strict family-level filtering (*q* = 1), where environmental conditions favor or disfavor entire families as units (see Appendix **??** for details). Competitive interactions were modeled via pairwise Ising couplings *J*_*ij*_, structured to reflect phylogenetic limiting similarity. We defined a contrast parameter *p* ∈ [0, 1] to modulate the specific cost of coexistence for closely related taxa. This parameter determines the divergence between within-family (*J*_*w*_) and between-family (*J*_*b*_) inhibitory strengths. Specifically, interactions are parameterized such that within-family inhibition scales as 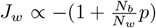, while between-family inhibition scales as *J*_*b*_ ∝ −(1 − *p*), where *N*_*w*_ and *N*_*b*_ are the counts of within- and between-family pairs, respectively. As *p* increases, the model imposes a severe penalty on the simultaneous high abundance of family members, enforcing competitive exclusion, whereas *p* = 0 implies uniform interactions independent of taxonomy.

## Acknowledgments

We thank M. Osella for inspiring the theoretical framework and M. Dal Bello, A. Goyal, L. Fant and M. Sireci for insightful discussions. O.M. acknowledges the Trieste Laboratory on Quantitative Sustainability - TLQS for funding. The work was funded by Fondo Italiano per la Scienza - FIS (CUP J53C23002290001; J.G.).

