## Supplementary text and figures for "A balance between environmental filtering and competitive exclusion modulates the macroecology of alternative stable states in microbial communities"

### Supplementary Information

#### A. Quantifying Bimodality

Although Figure 1B demonstrates clear bimodal patterns of distributions of relative abundances (AFDs) for representative ASVs, systematic analysis across all ASVs in our datasets required statistical metrics rather than visual assessment. Various moment-based metrics have been proposed to test bimodality [1], with the Sarle coefficient (based on skewness and kurtosis) being commonly used in recent studies [2–4]. An alternative approach is Hartigans' Dip Statistic, which measures deviations from unimodality using cumulative distribution properties rather than moments [5–7]. Given the known limitations of the Sarle coefficient and Hartigans' dip statistic [8, 9], we applied both methods, and, in addition we constructed another metric based on a likelihood ratio test between unimodal and bimodal model to quantify the bimodality of the AFD across ASVs.

##### A.1 Sarle Coefficient

Sarle coefficient, also known as the bimodal coefficient, is proportional to the ratio between squared skewness and kurtosis. For finite samples, the Sarle coefficient ($b$) is defined as [1]

$$b = \frac{g^2 + 1}{k + \frac{3(T-1)^2}{(T-2)(T-3)}} \tag{15}$$

where $T$ is the total samples, $g$ is the empirical skewness and $k$ is the empirical excess kurtosis. To quantify the bimodality of a given distribution, the Sarle coefficient is compared to a critical value of $5/9 \approx 0.555$ (the value for a uniform distribution), with values above this threshold indicating bimodality [9].

Figure SI.1 demonstrates that the majority of ASVs exhibit bimodal abundance distributions based on Sarle coefficient criteria. Computing the Sarle coefficient on relative abundance data, we observed that $75 - 100\%$ of ASVs are bimodal across all datasets, whether considering all ASVs or only those with high occupancy (occupancy 0.1). For multiple inocula experiments, the tendency for AFDs to be bimodal persists under log transformation irrespective of zero-handling method. In the multiple-inocula experiments, this tendency toward bimodality is robust to log transformation and to different zero-handling methods. In contrast, for the single-inoculum dataset we observe a markedly lower fraction of bimodal ASVs when applying a log transformation after adding one to read counts, or when replacing zero counts with 0.5 before computing relative abundance and taking the log. This shape occurs because most ASVs in this dataset exhibit bimodal patterns arising from the distinction between zero and non-zero reads. Since non-zero reads are often near detection limits, these transformations eliminate the meaningful separation between absent and present states, rendering the distributions unimodal. In contrast, replacing zeros with 0.01 preserves this separation and maintains higher bimodal detection rates.

##### A.2 Hartigans' Dip Statistic

The Hartigans' Dip Statistic is a measure of the dip (or the maximum difference) between the empirical cumulative distribution function and the cumulative distribution function of the best-fitting uniform distribution. Given the multiple modes in the multimodal distribution, it has a higher dip compared to a unimodal distribution. The Hartigan's Dip Statistic Python package [10] returns the dip statistic and its p-value. Thus, a distribution is identified as bimodal when its dip statistic is significantly larger than expected under a unimodal null hypothesis (p-value $< 0.05$). In the standard implementation, the p-value is computed by comparing against a null distribution generated assuming unimodality. However, we do not know whether this specific distribution is an appropriate null model for the purpose of this study.

Therefore, we developed a tailored approach using dip statistics with two null models: a unimodal model and a bimodal model, then compared the distribution of p-values across ASVs for each of the models. Unlike the standard approach that only tests against unimodality, our approach allows us to ask whether the AFD is unimodal or bimodal. In addition, having the distribution of the p-values gives us information on all ASVs rather than individual ASVs.

For the unimodal null model, we assumed that read count data is generated by a gamma distribution with Poisson sampling (negative binomial), while, for the bimodal model, we assumed that the data is generated by a two gamma distributions with Poisson sampling (two negative binomials) (see Section ). For each dataset, we fit read count data with the two models separately. Then, we generated 1,500 read count datasets from the fitted parameters of each null model. For each generated dataset, we computed the dip statistic for each ASV, then, calculated p-value as the proportion of null model dips smaller than the empirical dip.

In agreement with Sarle coefficient criteria, Hartigans' Dip Test results also indicate that majority of ASVs are bimodal. Figure SI.2 (top) shows the distribution of p-value of the relative abundance data across ASVs. For all dataset, we observed that there is a peak near zero for p-values against a unimodal null model. These peaks show that the majority of ASVs have an empirical dip that is larger than the model, from which we reject the unimodal hypothesis. On the other hand, the distribution of the p-values is uniform across ASVs with the mixture of gamma distribution model, indicating that the relative abundance of ASVs are bimodal. We also did the same computation for log-transformed dataset (Fig. SI.2). We handled the zeros by replacing them with 0.01 following the approach established in the Sarle coefficient analysis. The distribution of the p-values for the gamma model resembles that of the data without log transformation. For the bimodal model, the distribution of p-values is uniform for the single inoculum dataset, while it shows a bell-shaped distribution for the multiple inocula. Nevertheless, the absence of peak near zero for the bimodal model indicates that ASVs are bimodal.

#### A.3 Likelihood Ratio Test

While both the Sarle coefficient and dip test provide useful measures of bimodality, both have limitations. For the Sarle coefficient, it has an arbitrary critical value and potential misclassification of highly skewed distributions [8, 9]. For the dip test, while p-values indicate statistical significance, they do not quantify the degree that the bimodal model provides a better fit relative to the unimodal model. To address these limitations, we developed a more rigorous approach using likelihood ratio tests that directly compare the fit of unimodal and bimodal models from the data.

We performed a likelihood ratio test by computing the log of the likelihood ratio between the bimodal and unimodal models. For each ASV $i$, the log of the likelihood ratio ($\Delta\mathcal{L}_i$) is defined as

$$\Delta\mathcal{L}_i = \ln\left(\frac{\mathcal{L}_{b_i}}{\mathcal{L}_{u_i}}\right) = \ln\mathcal{L}_{b_i} - \ln\mathcal{L}_{u_i} \tag{16}$$

where $\mathcal{L}_{b_i}$ is the likelihood of the bimodal model (Eq. 6) with best fitting parameters and $\mathcal{L}_{u_i}$ is the likelihood of the unimodal model (Eq. 5) with best fitting parameters. Since a model with higher log-likelihood indicates a better fit to the observed data, we typically expect $\Delta\mathcal{L}_i$ for each ASV to be positive.

However, the positive trend of the log of the likelihood ratio can be trivial because a model with a greater degree of freedom provides a better fit compared to a more constrained model [11]. In our case, positive $\Delta\mathcal{L}_i$ value is expected since we compare a bimodal model (5 parameters) with a unimodal model (2 parameters). As described in the Section , the AFD follows a gamma distribution, but since our data consists of read counts, the sampling process results in a negative binomial distribution for read counts. The unimodal model (single gamma AFD + sampling or negative binomial) is a special case of the bimodal model (two-gamma mixture AFD + sampling or two negative binomial mixture). Because of that, we need to determine whether $\Delta\mathcal{L}_i$ is statistically significant.

We constructed a null model by generating read counts under the unimodal assumption and performed the same likelihood ratio test for each ASV. The reasoning is that if the AFD is truly bimodal then the empirical $\Delta\mathcal{L}_i$ will be larger than the typical log of likelihood ratio of the null model ($\langle\Delta\mathcal{L}_i\rangle_{\text{null}}$). In this case we generated data from the best fitted parameters of negative binomial distribution. We then computed $\Delta\mathcal{L}_{i,s}$ which we defined as the log of the likelihood ratio for ASV $i$ in a generated dataset $s$ and calculated

$$\langle\Delta\mathcal{L}_i\rangle_{\text{null}} = \frac{1}{S}\sum_{s=1}^{S}\Delta\mathcal{L}_{i,s}. \tag{17}$$

Figure SI.3 demonstrate that the majority of ASVs exhibit bimodal distributions. More than 75% of ASVs in each dataset (77% − 88%) have empirical log likelihood ratios significantly larger than expected under a unimodal null distribution. These percentages are consistent with our Sarle coefficient analysis of high occupancy ASVs, demonstrating that multiple independent statistical approaches converge on the same conclusion that the majority of ASVs are bimodal.

However, a small fraction of ASVs displayed different patterns. These ASVs have higher $\langle\Delta\mathcal{L}_i\rangle_{\text{null}}$ than the empirical values. In addition, there are cases of negative empirical values (3/30 in the single inoculum dataset and 3/108 in multiple inocula dataset with leucine), where the empirical value itself is less than $-10^{-3}$. It suggests that these ASVs might be unimodal rather than bimodal, but these ASVs are mostly rare ASVs.

In summary, we showed that the three bimodality metrics lead to the same conclusion: the majority of ASVs in each dataset are bimodal, despite their different underlying approaches (moment-based, cumulative distribution-based, and model-based). This convergence result across different method provides robust evidence that bimodality is a property of the Abundance Fluctuation Distributions in experimental microbial communities.

### B. Compositional Effects on Bimodal Abundance Distributions

A possible limitation of the bimodality analysis comes from the compositional nature of the data [12]. Since AFDs refer to relative abundances rather than absolute abundances across replicates, compositional effects may induce bimodality as an artifact of the transformation. For instance, if an ASV exhibits high variability in absolute abundance, it will dominate the total abundance in some replicates, forcing other ASVs to have low relative abundances (and vice versa), potentially creating artificial bimodal patterns. To investigate whether such compositional artifacts can occur and how to detect them, we developed a simple toy model to test whether one highly variable ASV could induce apparent bimodality in other ASVs. It must be highlighted that the aim of this toy model is to provide insight into compositional data behavior without any ecological relevance, and to serve as guidance for selecting the most appropriate data transformation method. Using this toy model, we evaluated common compositional transformation methods to determine which approach most effectively removes compositional artifacts. We then applied the most robust transformation to our empirical data to assess whether observed bimodality persists after eliminating the compositional effects.

We constructed a toy model consisting of five ASVs with mean true abundances ($\mu$): 300000, 175000, 47500, 25000, and 2500. We created scenarios in which one out of the five ASVs is bimodal by drawing the true abundances from mixture of Gaussian distributions with equal weight, with peaks located at $\mu \pm 0.2\mu$ and keeping the same CV for each peak of the mixture. The true abundances of the non-chosen ASVs were drawn from Gaussian distribution, centered at their respective mean values. For each of the scenario, we tested two different conditions based on the CV ratio between bimodal and unimodal ASVs ($r = \mathrm{CV}_{\mathrm{bimodal}}/\mathrm{CV}_{\mathrm{unimodal}}$). In the first condition, $r \approx 10$, while in the second condition $r \approx 1.0$.

Our first aim with the toy model was to check whether one bimodal ASV could introduce apparent bimodality in the unimodal ASVs when examining the relative abundance. Figure SI.4 displays the scenarios where ASV 0, ASV 2, or ASV 4 is bimodal when $r \approx 10$ (blue background) and where ASV 0 is bimodal with $r \approx 1.0$ (yellow background). For each scenario, we plotted the distribution of the centered-log of the true abundance (left) and the resulting relative abundance (right). When $r \approx 10$, the most abundant ASV, ASV 0, causes the apparent bimodal relative abundance of all of the other ASVs. As the mean of the bimodal ASV decreases, we observe the weakening of the apparent bimodality. For example, when ASV 4 (the rarest ASV) is bimodal, the relative abundance distributions of all ASVs retain the same unimodal/bimodal shape as their respective true abundances. In contrast, when $r \approx 1.0$, no bimodal distribution is observed in the relative abundance of unimodal ASVs, even the most abundant ASV is bimodal. Thus, our toy model suggests that apparent bimodality in relative abundance requires the bimodal ASV to have much higher CV than unimodal ASVs ($r \approx 10$). Moreover, the strength of this compositional effect increases with the mean abundance of the bimodal ASV.

Based on the toy model, we can check whether bimodal AFDs are the results of compositional effect by comparing the CV of true abundance of the highly abundant ASVs with the CV of rest of the ASVs. However, we do not have access to the true abundances. In order to find a good method that eliminates the compositionality effect, we tested two common and well-known transformations in this regard: the centered log-ratio (clr) and additive log-ratio (alr). In general, the two transformations remove the compositional effects by transforming the data from simplex to a real space [13].

Suppose that in a given sample $s$, there are $n$ ASVs with relative abundances $\mathbf{x}^s = (x_1^s \dots x_n^s)$ and geometric mean $g(\mathbf{x}^s) = \left(\prod_{i=1}^n x_i^s\right)^{1/n}$. The centered log-ratio (clr) transforms each of the relative abundances relative to the geometric mean of the sample as follows

$$\mathrm{clr}(\mathbf{x}^s) = \left(\log \frac{x_1^s}{g(\mathbf{x}^s)} \dots \log \frac{x_n^s}{g(\mathbf{x}^s)}\right). \tag{18}$$

This transformation is isometric and maintains the same number of ASVs in the transformed data. On the other hand, the additive log-ratio (alr) transforms the relative abundance relative to the relative abundance of an arbitrarily chosen ASV ($x_D$)

$$\mathrm{alr}(\mathbf{x}^s) = \left(\log \frac{x_1^s}{x_D^s}, \dots, \log \frac{x_{n-1}^s}{x_D^s}\right). \tag{19}$$

This transformation is not isometric, it transforms the data from $\mathcal{S}^n$ to $\mathbb{R}^{n-1}$.

Figure SI.5 shows the distributions of transformed relative abundances using three different methods: clr-transformation, alr-transformation with bimodal reference, and alr-transformation with unimodal reference. Here, we used the toy model where only ASV 0 is bimodal with $r \approx 10$ (Figure SI.4, top-left). Following clr-transformation, all ASVs are bimodal, indicating that this transformation cannot recover the underlying true abundance distributions. The alr-transformation with bimodal reference also cannot reveal the underlying true abundance distributions. In this case, the bimodal ASV (ASV 0) is the only possible choice as the bimodal reference, but when an ASV is used as its own reference, the transformation yields a delta-Dirac distribution (constant zero) for that ASV, which is trivial. Additionally, all other ASVs still have bimodal distributions. On the other hand, after alr-transformation with unimodal reference (ASV 1 as reference), the distribution of the transformed relative abundance of ASV 0 remains bimodal, while the other ASVs, except ASV 1, recover their unimodal distributions.

To explore the robustness of the alr-transformation with unimodal reference, we expand the model to have two bimodal ASVs. Figure SI.6 (top) shows the distribution of the true abundance and the distribution of the relative abundance of all ASVs when the true abundances of ASV 0 and ASV 2 are bimodal, while the true abundances of the other ASVs are unimodal. We noted that in this case, ASV 2 appears less bimodal in the relative abundance distribution because ASV 0 (the most abundant ASV) dominates the compositional effect, masking the pattern of the less abundant bimodal ASVs.

The results of each of the three transformation methods are still the same (Fig. SI.6 (bottom)). The clr-transformation still results in bimodal distributions for all ASVs. The reason is that any bimodal ASV makes the geometric mean of the relative abundances bimodal. It resulted into two clusters in the correlation between relative abundance of an ASV and the geometric mean. Each cluster correspond to one peak of the geometrical mean. Because of that, the clr-transformation preserves bimodal patterns rather than eliminating compositional artifacts. The alr-transformations with bimodal references result in the opposite pattern, while the same transformation with unimodal references can recover the true abundance pattern. The reason why the alr-transformation works is that when ASV 0 has high relative abundance, ASV 1 has low relative abundance, and vice versa. However, ASV 3 and ASV 1 both increase and decrease together. Dividing the relative abundance of ASV 0 by ASV 1 preserves the cluster structure because the opposing patterns create two distinct ratio values, maintaining the bimodal pattern. Conversely, dividing ASV 3 by ASV 1 produces similar ratio values because both vary proportionally, creating a single peak. Because of that, we can obtain back the shape of the true abundance distribution of ASV 0 (bimodal) and the shape of the true abundance distribution of ASV 3 (unimodal) in the alr-transformation with a unimodal reference.

The toy model shows us two important main points. First, one bimodal ASV can introduce apparent bimodality in the other ASVs if the CV of the bimodal ASV is much larger than the CV of the unimodal one and the mean of bimodal ASV is relatively high. Second, alr-transformation with unimodal reference is able to recover the bimodal/unimodal shape of the true abundances. The next step is to apply the alr-transformation to the data. While implementing the alr-method is straightforward, we were faced by the challenge of determining the reference. We do not have any prior information on the unimodality/bimodality of the true abundance.

Thus, instead of asking rigorous questions such as which ASVs are truly bimodal or which ASVs are bimodal due to compositional effect, we checked whether a single ASV induces apparent bimodality given its abundance. In particular, we tested the 5 most abundant ASVs in each dataset. If ASV $x$ was the only truly bimodal ASV among the 5, we would expect alr-transformation to reveal ASV $x$ as bimodal when referenced against any of the other 4 ASVs, while those 4 ASVs would appear unimodal when referenced against each other. Figure SI.7 - Figure SI.10 (top row) show the AFD of 5 most abundant ASVs in each data. Meanwhile, each column in the the $5 \times 5$ plots below it shows the alr-transformation of that ASV with any of the 5 ASVs as reference. In the single inoculum dataset, there is no case of unimodality in any ASV given any reference ASVs (Figure SI.7). Instead, we observed only bimodal or more complicated shape (multimodal). This holds under any possible bimodal ASV out of the 5 most abundant ASVs. The same is true for the multiple inocula dataset, where we also do not observe any case where exactly one ASV is bimodal while the other four are unimodal. We did notice some symmetric three-peak cases in the citrate dataset, but these represent additional complexity beyond simple bimodal/unimodal patterns. These results suggest that bimodal AFD in our datasets cannot be explained by simple compositional effects from a single bimodal ASV. The observed patterns more likely reflect either a true bimodality in multiple ASVs or more complex compositional interactions beyond what our toy model can cover.

To further investigate the compositional effect in the single inoculum dataset, we also employed the Optical Density (OD) data. Within a certain range, OD serves as a proxy for total cell counts [14, 15]. This allows us to estimate the absolute abundance of each ASV by multiplying its relative abundance by the OD of each sample. Figure SI.11 displays the estimated absolute abundance for five most abundant ASVs. Across all five ASVs and wavelengths, we observed two persistent peaks in the estimated true abundances. This finding, together with our previous results from alr-transformation of the data, confirms that the observed bimodality is likely not an artifact of the compositional nature of the data.

### C. Model Comparison: Gamma and Mixture of Two Gamma

In this section we compared the ability of the unimodal model (gamma) and the bimodal model (two gammas) to describe the AFDs. As shown in Section A.3, the majority of ASVs are bimodal, indicating that the two-gamma model better represents the AFD. To explore both models further, we generated 1,500 datasets from each model separately (see Section ). We compared three metrics: occupancy (fraction of samples where ASV is present), mean relative abundance, and coefficient of variation (CV). For each ASV, we computed each metric in every generated dataset, then averaged across the 1,500 datasets for comparison with empirical values.

Both models perform well for the mean relative abundance and occupancy (Figure SI.14). For the mean relative abundance, both models have very similar values but slightly overestimated mean relative abundance for low-abundance ASVs in the multiple inocula datasets. This overestimation may result from compositional effects during data generation, where for each generated dataset, the absence of rare ASVs in the process of generating the data increases the relative abundance of the other ASVs. Meanwhile for the occupancy, the two-gamma model shows less deviation from the empirical value compared to the gamma model.

In contrast, the two-gamma model outperforms the gamma model in reproducing the CV. The two-gamma model shows closer agreement with empirical CV values across all datasets. This indicates that the mixture model better captures the variance structure of abundance fluctuations compared to the single gamma distribution. We further explored the difference in the CV prediction of the two models by comparing the difference between the mean square error of the CV of the gamma model across all ASV and the mean square error of the CV of the two-gamma model across all ASV. We then constructed a null distribution by randomly shuffling the error between the two models independently for each ASV in each realization. For each realization, we computed the MSE of the each model across ASVs, then, we calculated the difference between the MSE of the two model (similar as the one with the data). The null distribution is the distribution of this difference across all realizations as shown in Figure SI.15. For multiple inocula dataset, we observed that the empirical differences (dashed line) lie on the right tail of the distribution, indicating that the CV predicted by the gamma has a significant larger error than the one from the two-gamma. On the other hand, for the single inoculum, the empirical difference does not lie near the tail, but at the 76th percentile of the null distribution. The smaller number of ASV in the single inoculum (30 ASVs) compared to the multiple inocula (80-110 ASVs) may be responsible for this difference. In addition, when we look into the empirical value of MSE of the two model, the MSE of the two-gamma model is approximately half of the MSE of the gamma model across all dataset (Figure SI.16). Thus, we conclude that the two-gamma model is better in explaining our data compared to the gamma model.

### D. Theoretical framework

Our empirical results suggest that alternative community states are organized around a small number of reproducible attractors, and that closely related ASVs exhibit reciprocal prevalence patterns *across* attractors but no systematic associations *within* attractors. To assess whether such structure can arise from a generic statistical mechanism, we develop a minimal Ising–Hopfield model with two components: (i) multi-attractor structure generated by Hopfield memories and (ii) competitive exclusion encoded by a block-structured interaction matrix.

#### D.1 Model definition

We represent each ASV $i$ by a binary variable

$$\sigma_i \in \{-1, +1\}, \tag{20}$$

indicating low- or high-abundance state respectively. A community configuration is

$$\boldsymbol{\sigma} = (\sigma_1, \ldots, \sigma_N) \ . \tag{21}$$

The probability of observing a given community configuration $P(\boldsymbol{\sigma})$ can be written, with no loss of generality, in the exponential form

$$P(\boldsymbol{\sigma}) = \frac{1}{Z} \exp\left(-\beta E(\boldsymbol{\sigma})\right) \ , \tag{22}$$

where the function $E$ (which in statistical physics corresponds to the energy) fully determines the joint probability distribution $P(\boldsymbol{\sigma})$. The parameter $\beta$ capture the effect of stochasticity: in the case $\beta = 0$ any configuration is equally probable (there is no effect of $E(\boldsymbol{\sigma})$). In the limit $\beta \to \infty$, only the configuration(s) minimizing the value of $E(\boldsymbol{\sigma})$ are permitted.

We assume that the function $E(\boldsymbol{\sigma})$ is defined as

$$E(\boldsymbol{\sigma}) = E_{\mathrm{mem}}(\boldsymbol{\sigma}) + E_{\mathrm{excl}}(\boldsymbol{\sigma}) \ , \tag{23}$$

where the two terms capture different mechanisms shaping community configurations. In particular, $E_{\mathrm{mem}}(\boldsymbol{\sigma})$ captures the alignment of the community configuration to an effective environmental state using a Hopfield memory term. This term may favor similar species to respond similarly to the same environmental state. On the other hand, $E_{\mathrm{excl}}(\boldsymbol{\sigma})$ may disfavor similar species to be in the same abundance state and captures exclusion and limiting similarity.

We assume that there exists $M$ traits matching with environmental states that favor or disfavor a given ASV $i$ to be in the high-abundance state. In particular, we define $M$ variables (memory pattern)

$$\boldsymbol{\xi}^\mu = (\xi_1^\mu, \ldots, \xi_N^\mu), \qquad \mu = 1, \ldots, M, \ , \tag{24}$$

and assume that

$$E_{\mathrm{mem}}(\boldsymbol{\sigma}) = -\frac{1}{2} \sum_\mu \left( \sum_j \xi_j^\mu \sigma_j \right)^2 \ . \tag{25}$$

Each of the $M$ patterns $\boldsymbol{\xi}^\mu$ defines a preferred configuration according to one "environmental axis" $\mu$. Community configurations will tend to align with the different environmental axes. Such alignment can be interpreted as the community state that ecological dynamics drive the system to. For instance, we can assume that ecological dynamics act on $M$ relevant ecological traits driving the system to community configurations with some specified functional profiles. The effect of the term (25) is to capture this ecological force. The variable $\xi_i^\mu$ therefore captures whether the ecological forces shaping the functional profile of an assembled community for a particular function/trait will tend to favor species $i$ being in high- or low-abundance.

In particular, we construct the $M$ patterns as randomly drawn binary variables

$$\xi_i^\mu \in \{-1, +1\} \ . \tag{26}$$

We expect similar species to have similar traits. We capture this effect by drawing these variables in a correlated manner. By building on the association between family and function [16], we assume that each memory $\mu$ has an associated "preferred family" $f^*(\mu)$, and the entries are drawn as

$$\mathbb{P}(\xi_i^\mu = +1) = \frac{1 + \sqrt{q}\, s_{f(i)}^\mu}{2}, \tag{27}$$

where $f(i)$ is the family of ASV $i$, and

$$s_{f(i)}^\mu = \begin{cases} 1, & f(i) = f^*(\mu), \\ \frac{1}{1-F}, & f(i) \neq f^*(\mu) \ , \end{cases} \tag{28}$$

where $F$ is the number of families. The parameter $q \in [0, 1]$ controls the degree of trait similarity / filtering

- $q = 0$: memories contain no family structure ("no filtering");
- $q = 1$: memories align strongly with families ("maximal filtering").

The term in eq. (25) can be rewritten as

$$E_{\mathrm{mem}}(\boldsymbol{\sigma}) = -\frac{1}{2} \sum_{i,j} J_{ij}^{\mathrm{mem}} \sigma_i \sigma_j \ , \tag{29}$$

where

$$J_{ij}^{\mathrm{mem}} = \sum_\mu \xi_i^\mu \xi_j^\mu \ . \tag{30}$$

To represent competitive exclusion we assume

$$E_{\mathrm{excl}}(\boldsymbol{\sigma}) = -\frac{1}{2} \sum_{i \neq j} J_{ij}^{\mathrm{excl}} \sigma_i \sigma_j \ . \tag{31}$$

Since we want to model the possibility that competition and exclusion are stronger among closely related taxa, we introduce a family block structure:

$$J_{ij}^{\mathrm{excl}} = \begin{cases} J_w, & f(i) = f(j),\ i \neq j, \\ J_b, & f(i) \neq f(j), \end{cases} \qquad J_w \leq J_b \leq 0. \tag{32}$$

The magnitudes $J_w, J_b$ are chosen such that the total sum $\sum_{i<j} J_{ij}^{\mathrm{excl}} = -N J_{\mathrm{mean}}$ is fixed, ensuring that exclusion is redistributed among? within- and between-family pairs rather than increased overall when the relationship between $J_w$ and $J_b$ changes.

When all $F$ families contain the same number of species $n = N/F$, the number of unordered pairs within and between families can be written explicitly. Within-family pairs:

$$N_w \ = \ F \binom{n}{2} \ = \ \frac{F\, n(n-1)}{2}, \tag{33}$$

and between-family pairs:

$$N_b \;=\; \binom{N}{2} - N_w \;=\; \frac{N(N-1)}{2} - \frac{F\,n(n-1)}{2}. \tag{34}$$

Using $N = Fn$, one obtains the ratio

$$\frac{N_b}{N_w} = \frac{Fn - n - (n-1)}{n-1} = \frac{n(F-1)}{n-1}. \tag{35}$$

With the parametrization

$$J_w = -J_{\text{mean}}(1 + \frac{N_b}{N_w}p), \qquad J_b = -J_{\text{mean}}\,(1-p)\,, \tag{36}$$

we introduce the contrast parameter $p \in [0,1]$ that controls the strength of within-family exclusion: high contrast means $J_w$ is much more negative than $J_b$.

The difference between within- and between-family couplings becomes

$$J_w - J_b = -pJ_{\text{mean}}\left(1 + \frac{N_b}{N_w}\right) = -p\,J_{\text{mean}}\left(1 + \frac{n(F-1)}{n-1}\right). \tag{37}$$

Thus, in the equal-sized case the contrast parameter $p$ controls the coupling gap linearly:

$$|J_w - J_b| = p\,J_{\text{mean}}\left(1 + \frac{n(F-1)}{n-1}\right). \tag{38}$$

In particular, for large families ($n \gg 1$) this simplifies to

$$|J_w - J_b| \approx p\,J_{\text{mean}}F. \tag{39}$$

When $p > 0$, the form of eq. 31 strongly penalizes configurations in which many members of the same family take the same state.

Combining both contributions, the total Hamiltonian is

$$E(\boldsymbol{\sigma}) = -\frac{1}{2}\sum_{i\neq j}\left[J_{ij}^{\text{mem}} \;+\; J_{ij}^{\text{excl}}\right]\sigma_i\sigma_j\ . \tag{40}$$

#### D.2 Family-level mean-field free energy and analytical transition line

To analyze how the memory strength, exclusion penalty, and environmental filtering combine to shape attractor structure, we derive a mean-field free energy directly in terms of family-resolved order parameters.

We recall that we partition the $N$ ASVs into $F$ equal-sized families of size $n = N/F$, and define:

$$m_f \;=\; \frac{1}{n}\sum_{i\in f}\sigma_i, \qquad S_f = n\,m_f, \tag{41}$$

and the family-level overlap with memory $\mu$

$$m_{\mu,f} = \frac{1}{n}\sum_{i\in f}\xi_i^{\mu}\sigma_i. \tag{42}$$

Patterns are drawn with a family-level bias controlled by the filtering parameter $q$:

$$\mathbb{P}(\xi_i^{\mu} = +1 \mid f(i) = f) = \frac{1 + \sqrt{q}\,s_f^{\mu}}{2}, \qquad \langle \xi_i^{\mu}\rangle_f = \sqrt{q}\,s_f^{\mu}, \tag{43}$$

where $s_f^{\mu} = 1$ for the family preferred by pattern $\mu$, and $s_f^{\mu} = -1/(F-1)$ otherwise.

Now we make a *mean field approximation within families* and write

$$m_{\mu,f} = \frac{1}{n}\sum_{i\in f}\xi_i^{\mu}\sigma_i \approx \langle \xi_i^{\mu}\rangle_f\, m_f = \sqrt{q}\,s_f^{\mu}\,m_f. \tag{44}$$

Thus the filtering parameter $q$ enters through the replacement

$$m_{\mu,f} \approx \sqrt{q}\,s_f^{\mu}m_f, \qquad m_{\mu} = \frac{1}{F}\sum_f m_{\mu,f} \approx \frac{\sqrt{q}}{F}\sum_f s_f^{\mu}m_f. \tag{45}$$

For $q = 1$ and $m_f = s_f^\mu$ this expression for $m_\mu$ saturates at $m_\mu = 1/(F-1)$ instead of 1. We introduce a renormalization factor chosen so that the approximate expression attains the correct saturation value:

$$m_\mu \approx \frac{\sqrt{q}(F-1)}{F} \sum_f s_f^\mu m_f. \tag{46}$$

For a single dominant memory $\mu^\star$ (retrieval state), the Hopfield energy becomes

$$E_{\text{mem}} = -\frac{1}{2}\left(\sum_i \xi_i^{\mu^\star} \sigma_i\right)^2 \approx -q\frac{n^2(F-1)^2}{2}\left(\sum_f s_f^{\mu^\star} m_f\right)^2. \tag{47}$$

The exclusion energy decomposes exactly as

$$E_{\text{excl}} = \frac{|J_w - J_b|}{2}\sum_f S_f^2 \; + \; (\text{global and constant terms}) = \frac{|J_w - J_b|}{2} n^2 \sum_f m_f^2 \; + \; \text{const}. \tag{48}$$

Therefore, by using eq.(39) for $n \gg 1$, we can write

$$E_{\text{excl}} \approx p\frac{J_{\text{mean}} F n^2}{2}\sum_f m_f^2. \tag{49}$$

In the low-stochasticity regime ($\beta \to \infty$), entropic contributions are negligible, and the relevant mean-field description is obtained directly from the coarse-grained energy expressed in terms of the family magnetizations. Thus, the low-stochasticity coarse energy is the quadratic form

$$E(\mathbf{m}) = \frac{1}{2}\mathbf{m}^\top \left(A\,I \; - \; B\,\mathbf{s}\mathbf{s}^\top\right)\mathbf{m}, \qquad A \equiv pJ_{\text{mean}} F n^2, \quad B \equiv qn^2(F-1)^2, \tag{50}$$

where $\mathbf{s} = (s_1^{\mu^\star}, \ldots, s_F^{\mu^\star})$.

All directions orthogonal to $\mathbf{s}$ have eigenvalue $A > 0$ and are therefore stable. Along $\mathbf{s}$ itself,

$$\lambda_\parallel = A - B\,\|\mathbf{s}\|^2. \tag{51}$$

The symmetric state $m_f = 0$ becomes unstable precisely when $\lambda_\parallel < 0$, i.e.

$$A < B\,\|\mathbf{s}\|^2. \tag{52}$$

For the biased family structure used here,

$$\|\mathbf{s}\|^2 = 1 + \frac{1}{F-1} = \frac{F}{F-1}, \tag{53}$$

and the instability condition becomes

$$q > p\,\frac{J_{\text{mean}}}{F-1}. \tag{54}$$

Thus, in the low-stochasticity limit, the transition from an *exclusion-dominated* regime (families unable to magnetize coherently) to a *filtering-dominated* regime (families aligning along memory patterns), happens at the threshold

$$q \; = \; p\,, \tag{55}$$

if we set $J_{\text{mean}}/(F-1) = 1$.

#### D.3 Simulations

We simulate dynamics of the Ising community using single-spin Metropolis updates at low stochasticity $\beta \gg 1$. At each update, a spin $i$ is selected uniformly at random and a flip $\sigma_i \to -\sigma_i$ is proposed. The energy change reads

$$\Delta E_i = 2\sigma_i \sum_j J_{ij}\sigma_j, \qquad J_{ij} = J_{ij}^{\text{mem}} + J_{ij}^{\text{excl}}, \tag{56}$$

and the flip is accepted with probability $\min(1, e^{-\beta \Delta E_i})$. A simulation consists of $N_{\text{sweeps}}$ sweeps, where each sweep corresponds to $N$ such attempted updates. Different assembly histories are represented by independent trajectories initialized from independent random spin configurations.

To enable a direct comparison between model simulations and empirical data, we define attractors, ASV prevalence, and pairwise prevalence relationships using the same notation and criteria as in the data analysis. For a fixed parameter pair $(q, p)$, we run $N_{\text{runs}}$ independent assembly trajectories starting from random initial conditions. Terminal community configurations are clustered in configuration space into $M$ clusters using $k$-means, which we interpret as model attractors and label by $\alpha = 1, \ldots, M$. Let $N_\alpha$ denote the number of simulation runs assigned to attractor $\alpha$.

For each ASV $i$ and attractor $\alpha$, we define the prevalence

$$\pi_{i,\alpha} = \frac{1}{N_\alpha} \sum_{r \in \alpha} \mathbb{I}\left(\sigma_i^{(r)} = +1\right), \tag{57}$$

where $\sigma_i^{(r)}$ is the terminal abundance state of ASV $i$ in simulation run $r$, and $\mathbb{I}(\cdot)$ is the indicator function. As in the data, $\pi_{i,\alpha}$ represents the probability that ASV $i$ is in the high-abundance state within attractor $\alpha$.

Prevalence values are discretized into high- and low-prevalence categories using the same threshold as in the empirical analysis. We define the binary prevalence-pattern matrix

$$\mathcal{B}_{i,\alpha} = \mathbb{I}\left(\pi_{i,\alpha} \geq \frac{1}{2}\right), \tag{58}$$

so that each ASV $i$ is associated with a binary vector $\mathbf{b}_i = (\mathcal{B}_{i,1}, \ldots, \mathcal{B}_{i,M})$ describing its prevalence pattern across attractors.

For each pair of ASVs $(i, j)$ belonging to the same family, we compare their binary prevalence patterns $\mathbf{b}_i$ and $\mathbf{b}_j$:

- The pair is classified as *same* if $\mathbf{b}_i = \mathbf{b}_j$, i.e. the two ASVs share identical prevalence classifications across all attractors.
- The pair is classified as *reciprocal* if $\mathbf{b}_i = \mathbf{1} - \mathbf{b}_j$, such that one ASV is high-prevalence precisely where the other is low-prevalence.
- All remaining pairs are classified as *different*.

Let $N^{\text{in}}_{\text{same}}$, $N^{\text{in}}_{\text{recip}}$, and $N^{\text{in}}_{\text{diff}}$ denote the numbers of within-family ASV pairs in each category. We report the within-family fractions

$$f_{\text{same}} = \frac{N^{\text{in}}_{\text{same}}}{N^{\text{in}}_{\text{same}} + N^{\text{in}}_{\text{recip}} + N^{\text{in}}_{\text{diff}}}, \qquad f_{\text{recip}} = \frac{N^{\text{in}}_{\text{recip}}}{N^{\text{in}}_{\text{same}} + N^{\text{in}}_{\text{recip}} + N^{\text{in}}_{\text{diff}}}. \tag{59}$$

To assess whether the observed within-family fractions are larger than expected by chance taxonomy, we use a null model that *preserves* the prevalence-pattern matrix $\mathcal{B}$ but *shuffles* family assignments across ASVs (as in the empirical analysis). For each shuffle $s = 1, \ldots, N_{\text{null}}$, we recompute the within-family fractions $f^{\text{null}}_{\text{same},s}$ and $f^{\text{null}}_{\text{recip},s}$, and define the null means

$$\bar{f}^{\text{null}}_{\text{same}} = \frac{1}{N_{\text{null}}} \sum_{s=1}^{N_{\text{null}}} f^{\text{null}}_{\text{same},s}, \qquad \bar{f}^{\text{null}}_{\text{recip}} = \frac{1}{N_{\text{null}}} \sum_{s=1}^{N_{\text{null}}} f^{\text{null}}_{\text{recip},s}. \tag{60}$$

We summarize deviations from the null with enrichment ratios

$$R_{\text{same}} = \frac{f_{\text{same}}}{\bar{f}^{\text{null}}_{\text{same}}}, \qquad R_{\text{recip}} = \frac{f_{\text{recip}}}{\bar{f}^{\text{null}}_{\text{recip}}}. \tag{61}$$

Unless otherwise stated, simulations use $N = 40$ ASVs, $F = 4$ families, $M = 4$ memories/attractors, $J_{\text{mean}} = F - 1$, $\beta = 100$, $N_{\text{sweeps}} = 1000$, and $N_{\text{runs}} = 200$. We show in Fig. SI.34 A an example of a prevalence pattern for $q = 0.5$ and $p = 0.9$. We scan a grid of parameters $p \in [0, 1]$ (exclusion contrast) and $q \in [0, 1]$ (filtering strength in the memory construction), and for each point we average results over $K$ independent realizations (independent memories, couplings, and dynamics). We report the mean across $K = 20$ realizations for $R_{\text{same}}$ and $R_{\text{recip}}$, producing phase-diagram heatmaps for both observables (Fig. SI.34 B).

For all combinations of $q$ and $p$ taking values 0.1, 0.5, and 0.9, representative of low (L), medium (M), and high (H) importance, we compare model outcomes against the matched null model using boxplots. For each of $K = 200$ independent realizations we store both the simulation fractions $(f^{(k)}_{\text{same}}, f^{(k)}_{\text{recip}})$ and the corresponding null means $(\bar{f}^{\text{null},(k)}_{\text{same}}, \bar{f}^{\text{null},(k)}_{\text{recip}})$ computed from shuffles of the same $B^{(k)}$. This yields paired distributions over $k = 1, \ldots, K$ for both simulation and null quantities, enabling direct hypothesis tests. In particular, we compute two-sided permutation p-values, with $10^4$ permutations per comparison, by comparing the distributions of empirical and null means across realizations; significance is summarized with asterisks, with $*$ ($< 0.05$), $**$ ($< 0.01$), $***$ ($< 0.001$) (Fig. SI.34 C).

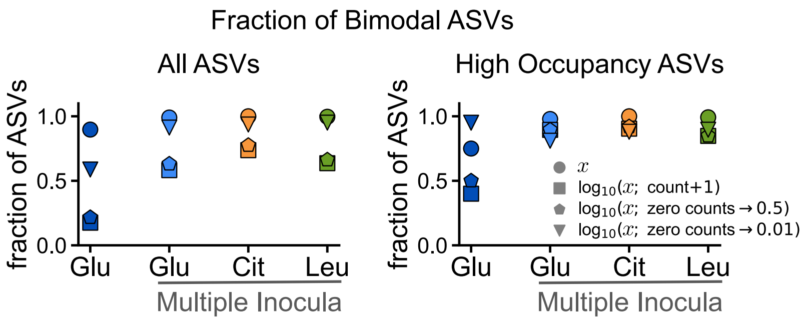


**Figure SI.1: Majority of ASVs exhibit bimodal abundance distributions based on Sarle coefficient criterion.** Fraction of bimodal ASVs (Sarle coefficient $> 5/9 \approx 0.55$) for all ASVs (left) and ASVs with occupancy $> 0.1$ (right) for all experimental datasets. Marker shapes denote the data transformation used to compute the Sarle coefficient with relative abundance, $x =$ counts/total counts.

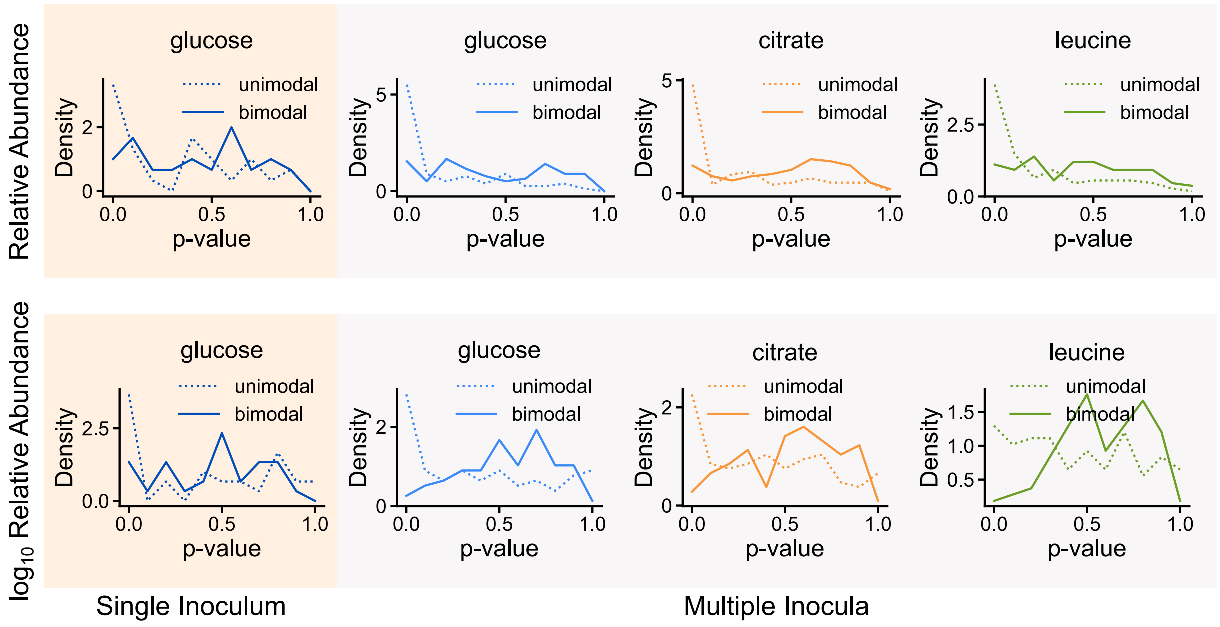


**Figure SI.2: Hartigan's Dip Test supports bimodality of ASV distributions.** Distributions of p-values for unimodal (dotted) vs bimodal (solid) null hypotheses using relative abundance (top) and $\log_{10}$-transformed data with zero counts replaced by 0.01 (bottom). Peaks near zero for unimodal hypothesis indicate rejection of unimodality.

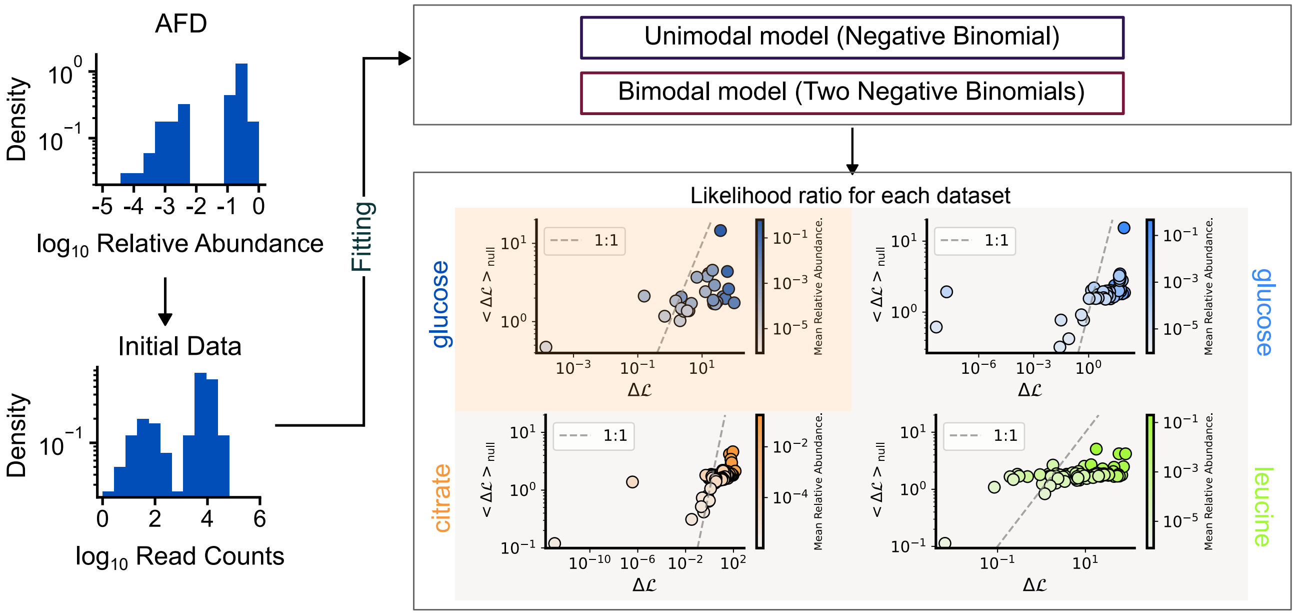


**Figure SI.3: A log-likelihood ratio method demonstrates that empirical AFDs are bimodal.** Bimodality is quantified by comparing unimodal and bimodal model fits (Equation 16). Scatter plots show the log-likelihood ratio ($\Delta\mathcal{L}$) is significantly larger for empirical data than for a unimodal null model, confirming widespread bimodality.

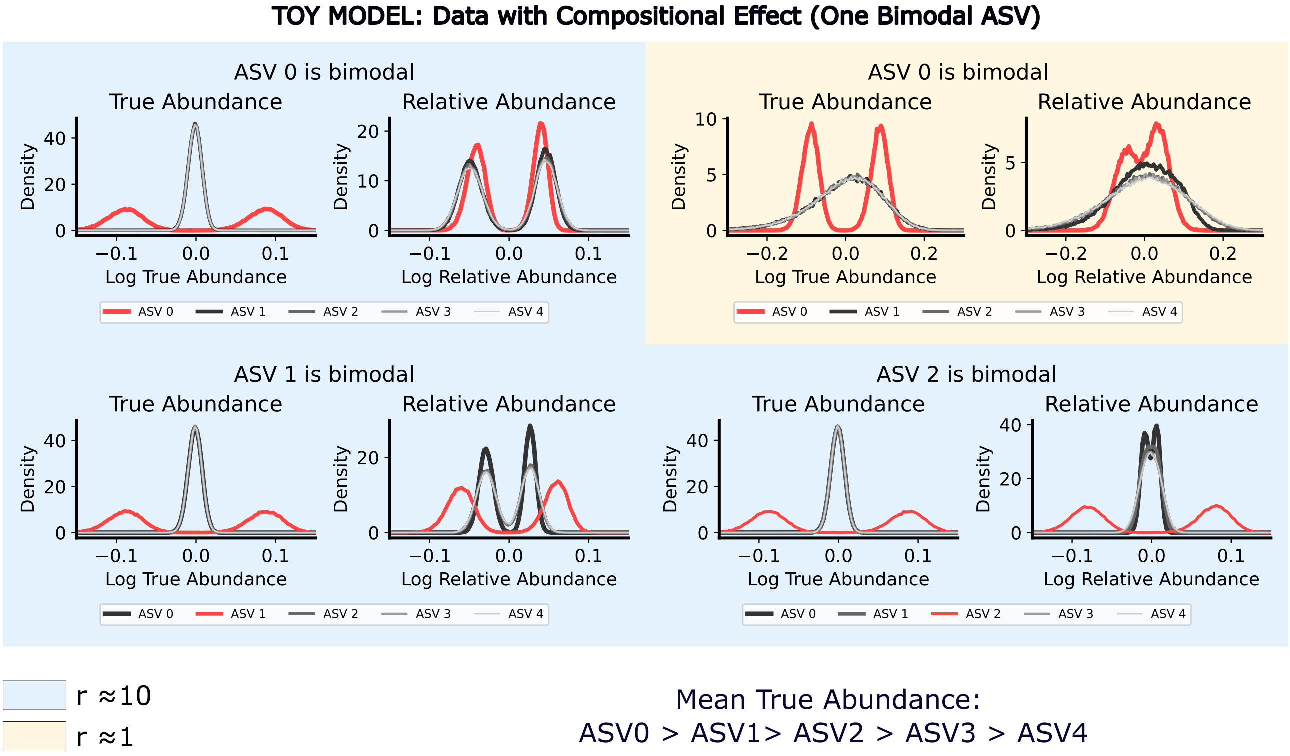


**Figure SI.4: A single bimodal ASV induces compositional effects only when CV ratios are large.** Toy model with 5 ASVs where only one ASV is truly bimodal (shown in red) while others are unimodal, with decreasing mean abundances (ASV 0 > ASV 1 > ASV 2 > ASV 3 > ASV 4). Four scenarios test different ASVs as the bimodal one: ASV 0 with $r \approx 10$ (top left), ASV 0 with $r \approx 1$ (top right), ASV 1 with $r \approx 10$ (bottom left), and ASV 2 with $r \approx 10$ (bottom right). Each scenario shows true abundance distributions (left) and resulting relative abundance distributions (right). When $r \approx 10$ ($\mathrm{CV}_{\mathrm{bimodal}} \gg \mathrm{CV}_{\mathrm{unimodal}}$), the single bimodal ASV creates apparent bimodality in unimodal ASVs' relative abundances, with strongest effects when high-abundance ASVs are bimodal. When $r \approx 1$, no compositional effects occur.

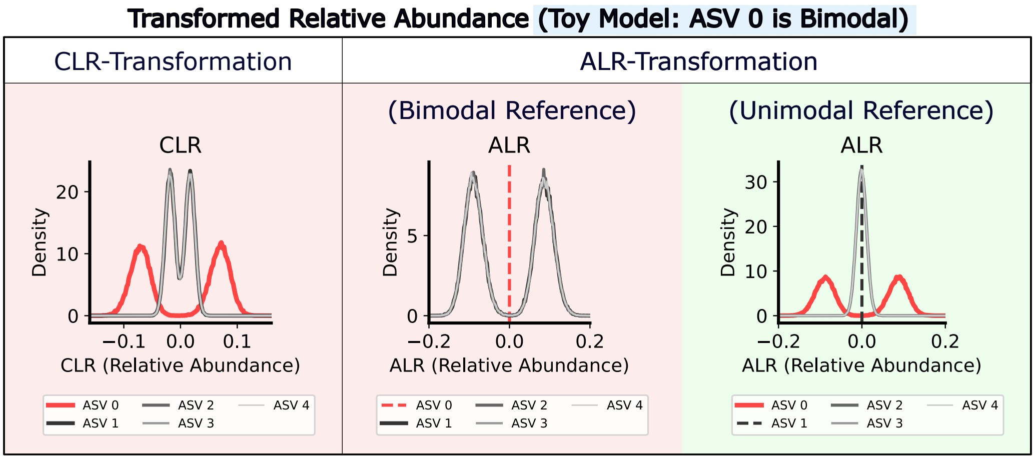


**Figure SI.5: Alr transformation with unimodal reference recovers true abundance distributions.** Three transformations applied to the toy model case where ASV 0 is bimodal (top-left of Figure SI.4): centered log-ratio (clr) transformation (left), additive log-ratio (alr) transformation with bimodal reference (middle), and alr transformation with unimodal reference (right). clr transformation fails to remove compositional effects, maintaining apparent bimodality in all ASVs. alr with bimodal reference gives trivial results for the reference ASV and maintains apparent bimodality in others. alr with unimodal reference (ASV 1) successfully recovers the original unimodal distributions while preserving the true bimodal pattern in ASV 0.

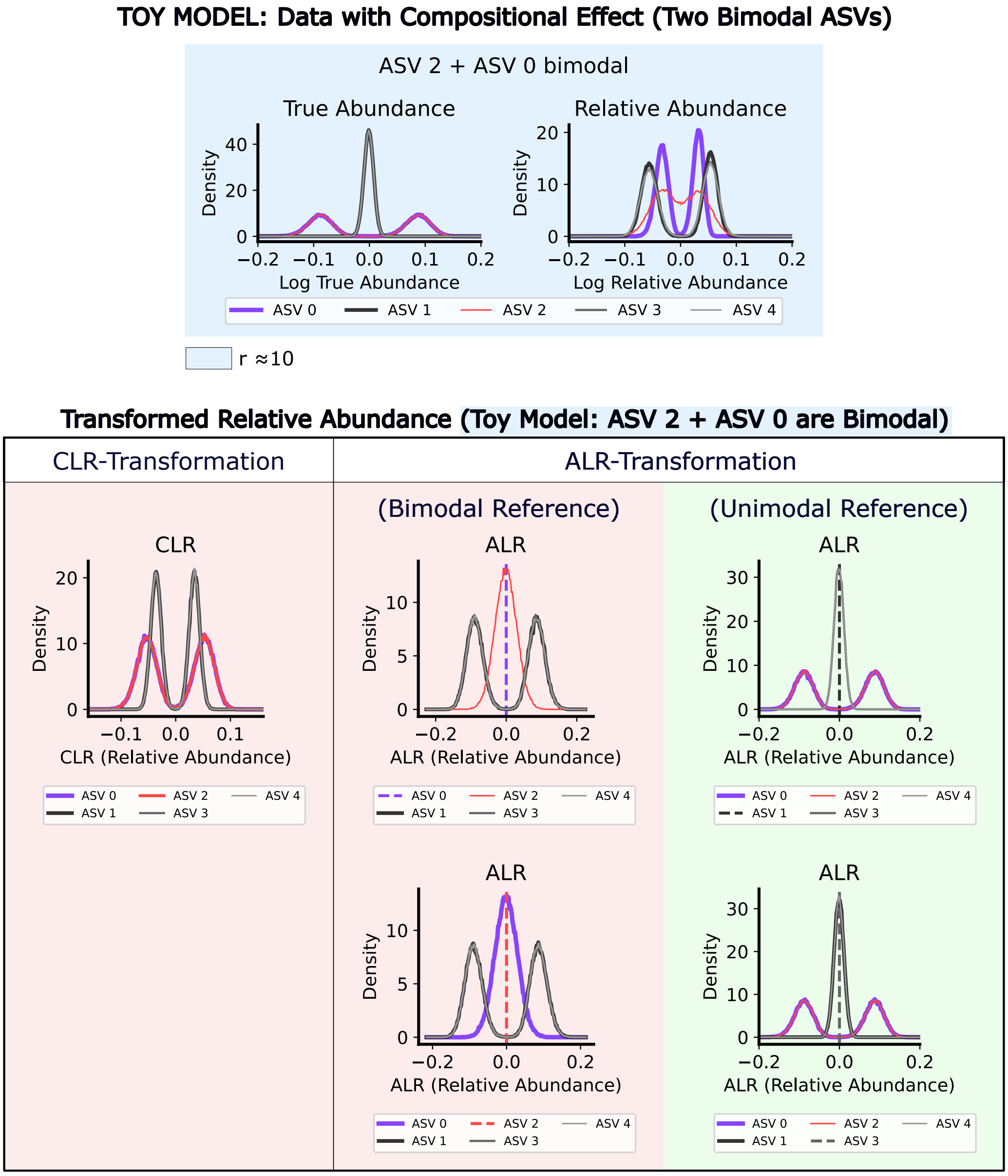


**Figure SI.6: Alr transformation with unimodal reference recovers true abundance distributions even with multiple bimodal ASVs.** Toy model with ASV 0 and ASV 2 as bimodal ASVs ($r \approx 10$) while others remain unimodal. Two bimodal ASVs create more complex apparent bimodality in relative abundances (top). The same three transformations (bottom) show that only alr transformation with unimodal reference successfully removes compositional effects.

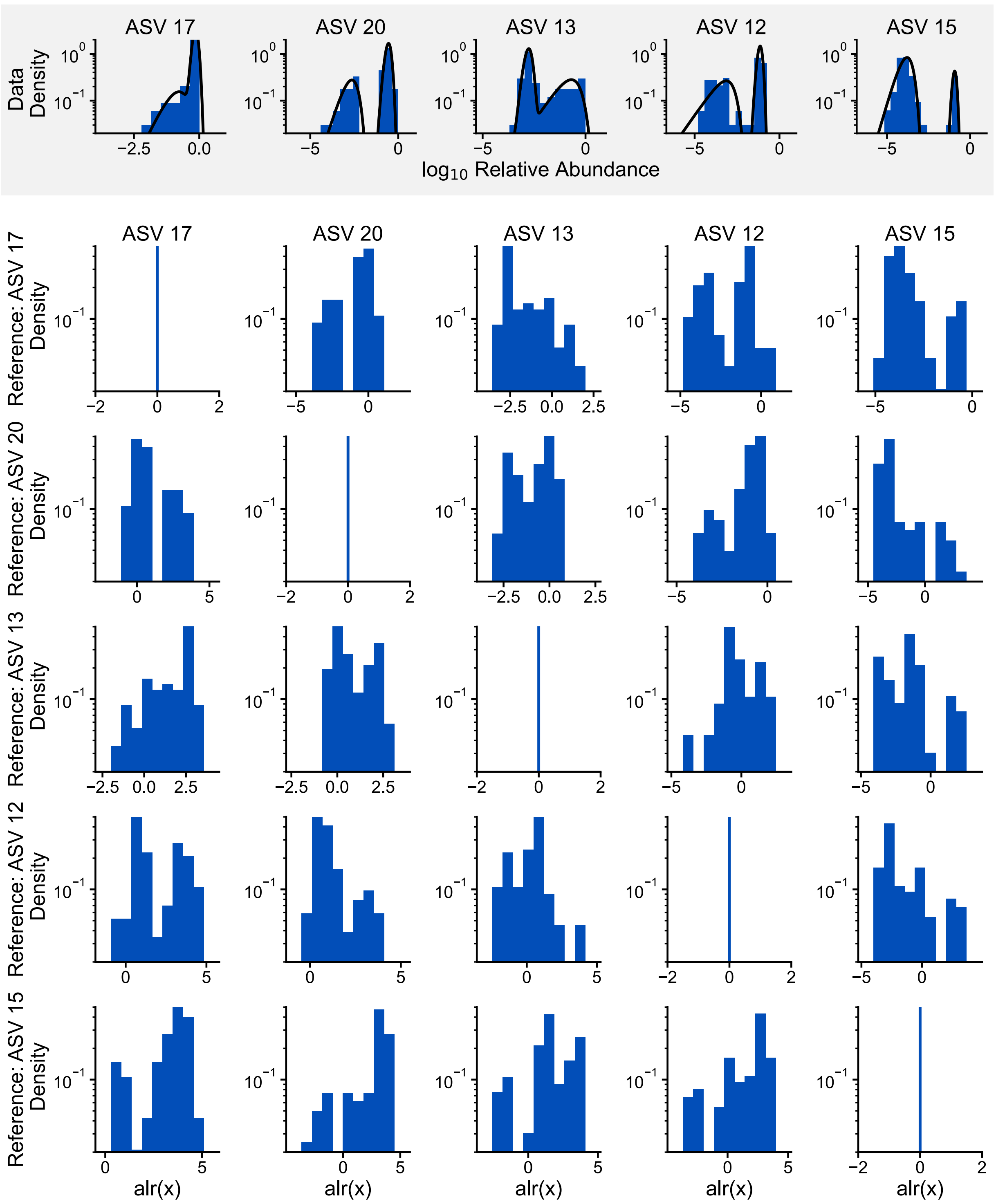


**Figure SI.7: No single ASV induces apparent bimodality in glucose (single inoculum).** Top row (grey background) shows AFDs of the 5 ASVs with highest average abundance across samples (ASV 17, ASV 20, ASV 13, ASV 12, ASV 15; left to right: highest to lowest average abundance). The 5×5 matrix below shows alr-transformed relative abundances where each column corresponds to one target ASV (ASV 17, ASV 20, ASV 13, ASV 12, ASV 15 from left to right) and each row uses a different reference ASV (ASV 17, ASV 20, ASV 13, ASV 12, ASV 15 from top to bottom). Diagonal panels show trivial cases where each ASV serves as its own reference.

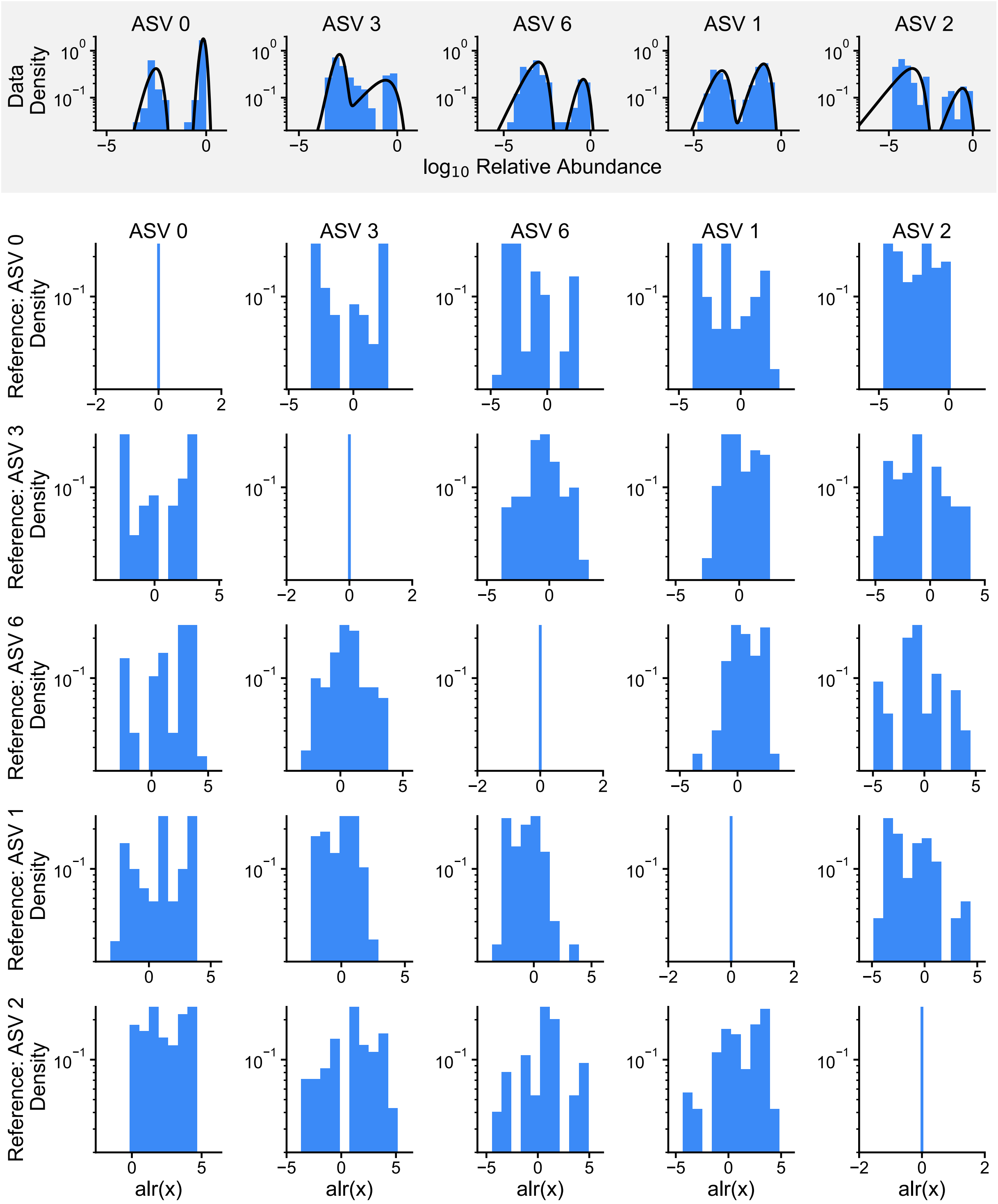


**Figure SI.8: No single ASV induces apparent bimodality in glucose (multiple inocula).** Top row (grey background) shows AFDs of the 5 ASVs with highest average abundance across samples (ASV 17, ASV 20, ASV 13, ASV 12, ASV 15; left to right: highest to lowest average abundance). The 5×5 matrix below shows alr-transformed relative abundances where each column corresponds to one target ASV (ASV 17, ASV 20, ASV 13, ASV 12, ASV 15 from left to right) and each row uses a different reference ASV (ASV 17, ASV 20, ASV 13, ASV 12, ASV 15 from top to bottom). Diagonal panels show trivial cases where each ASV serves as its own reference.

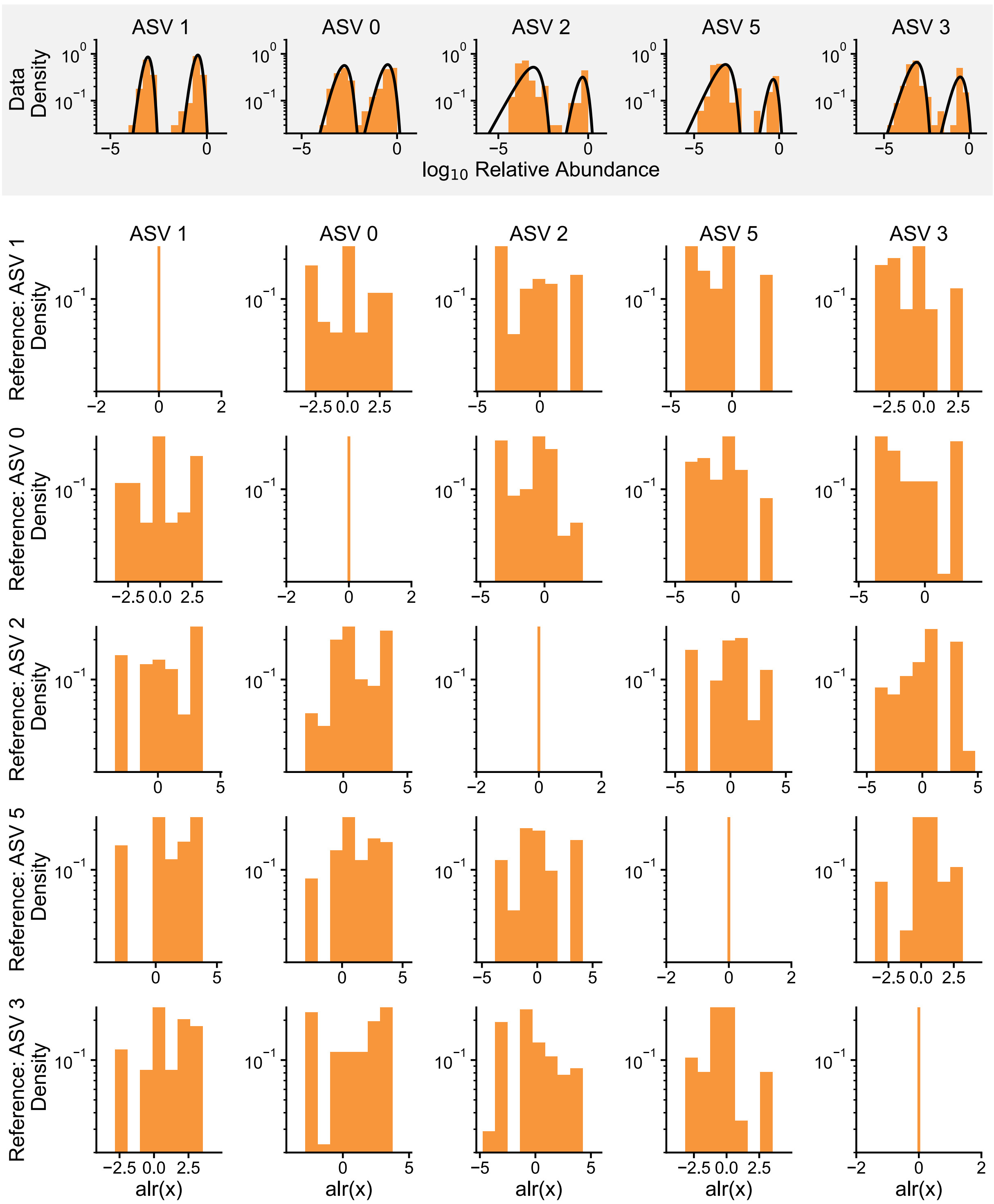


**Figure SI.9: No single ASV induces apparent bimodality in citrate (multiple inocula).** Top row (grey background) shows AFDs of the 5 ASVs with highest average abundance across samples (ASV 17, ASV 20, ASV 13, ASV 12, ASV 15; left to right: highest to lowest average abundance). The 5×5 matrix below shows alr-transformed relative abundances where each column corresponds to one target ASV (ASV 17, ASV 20, ASV 13, ASV 12, ASV 15 from left to right) and each row uses a different reference ASV (ASV 17, ASV 20, ASV 13, ASV 12, ASV 15 from top to bottom). Diagonal panels show trivial cases where each ASV serves as its own reference.

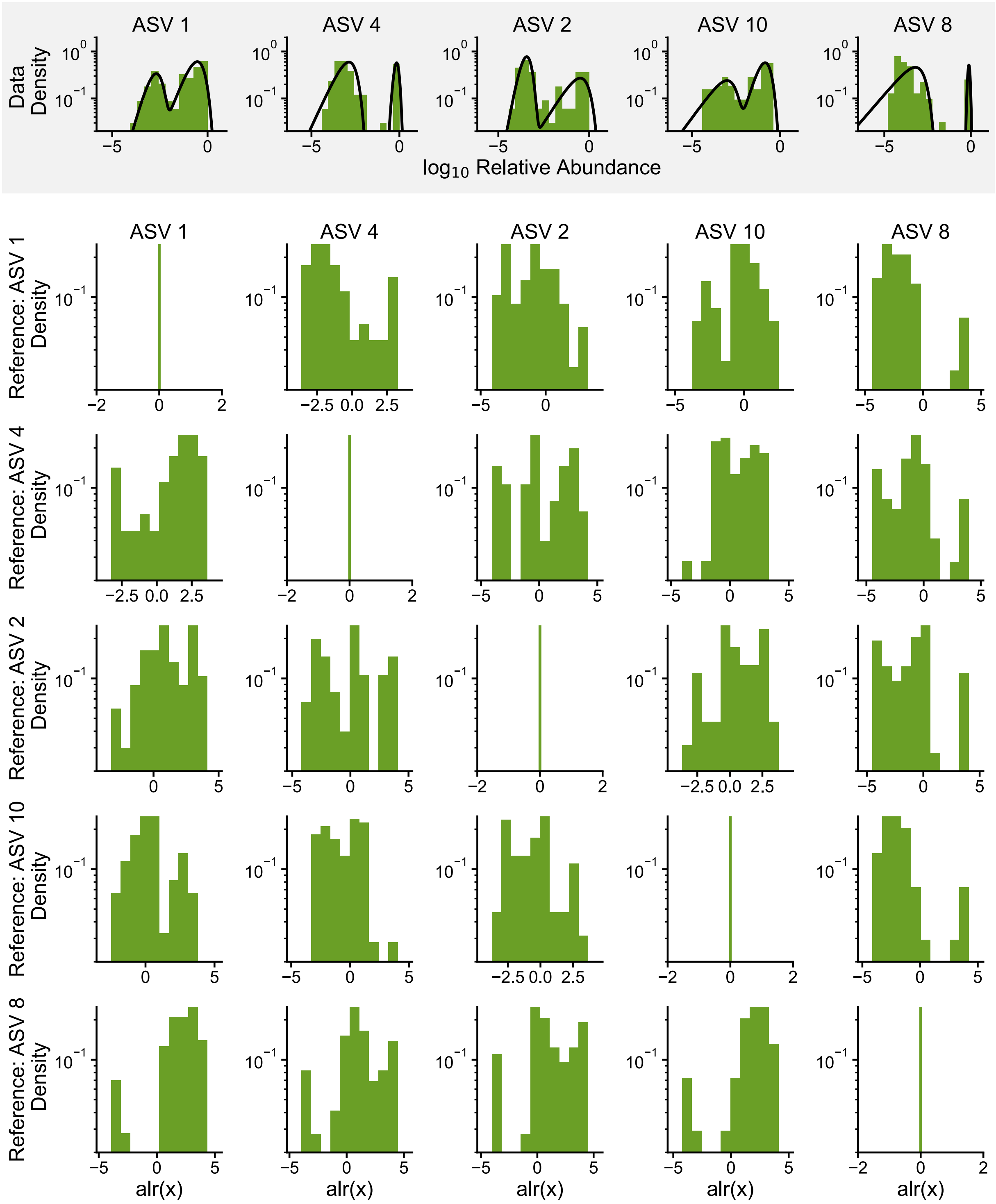


**Figure SI.10: No single ASV induces apparent bimodality in leucine (multiple inocula).** Top row (grey background) shows AFDs of the 5 ASVs with highest average abundance across samples (ASV 17, ASV 20, ASV 13, ASV 12, ASV 15; left to right: highest to lowest average abundance). The 5×5 matrix below shows alr-transformed relative abundances where each column corresponds to one target ASV (ASV 17, ASV 20, ASV 13, ASV 12, ASV 15 from left to right) and each row uses a different reference ASV (ASV 17, ASV 20, ASV 13, ASV 12, ASV 15 from top to bottom). Diagonal panels show trivial cases where each ASV serves as its own reference.

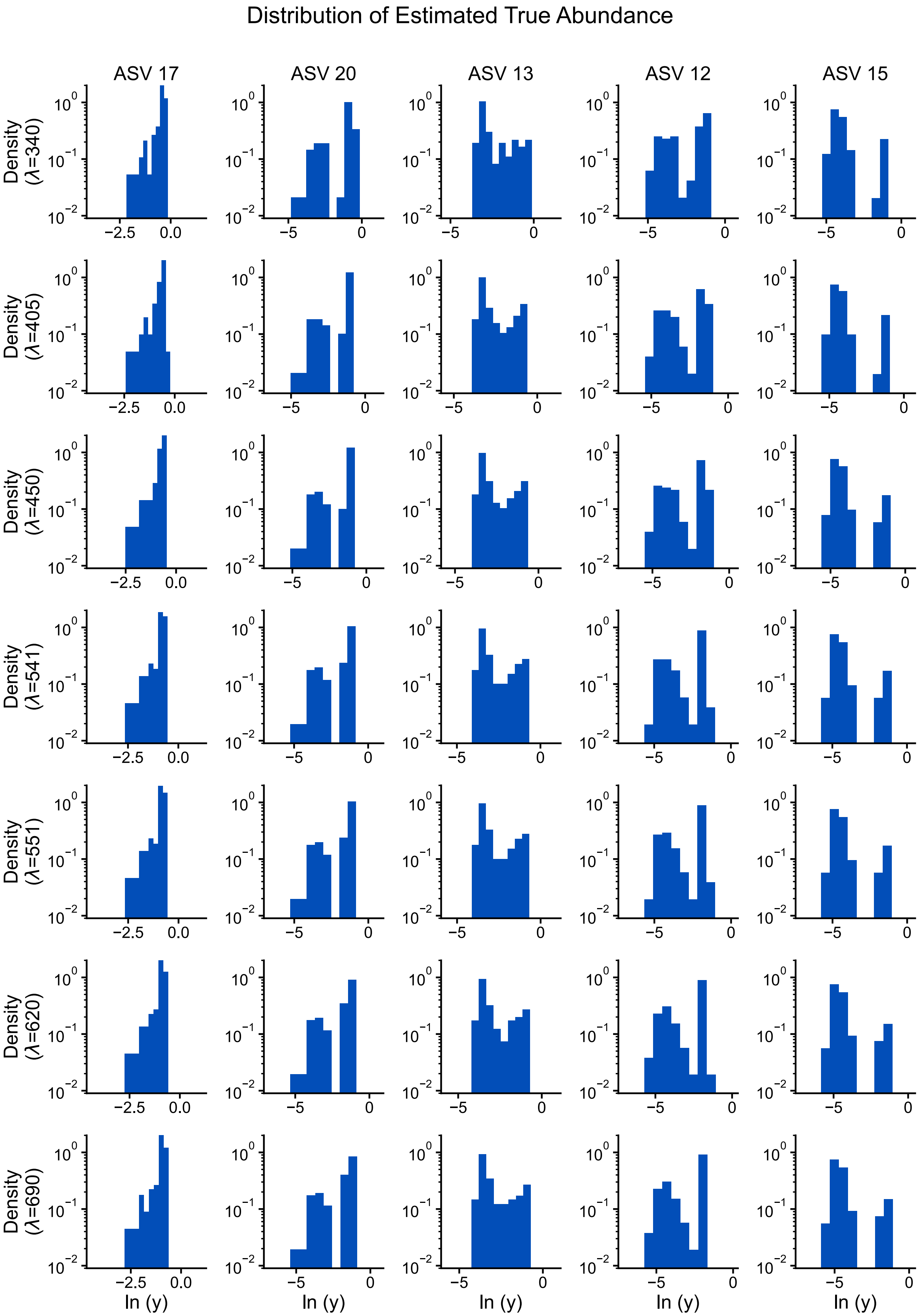


**Figure SI.11: Bimodality persists in estimated absolute abundances across all OD wavelengths in glucose (single inoculum).** Distribution of estimated absolute abundances (y = relative abundance × OD) for the 5 ASVs with highest average abundance (ASV 17, ASV 20, ASV 13, ASV 12, ASV 15; left to right: highest to lowest average abundance). Each row shows results for different OD wavelengths (λ=340, 405, 450, 541, 551, 620, 690 nm from top to bottom). Persistent bimodal patterns across all ASVs and wavelengths confirm that observed bimodality is not a compositional artifact.

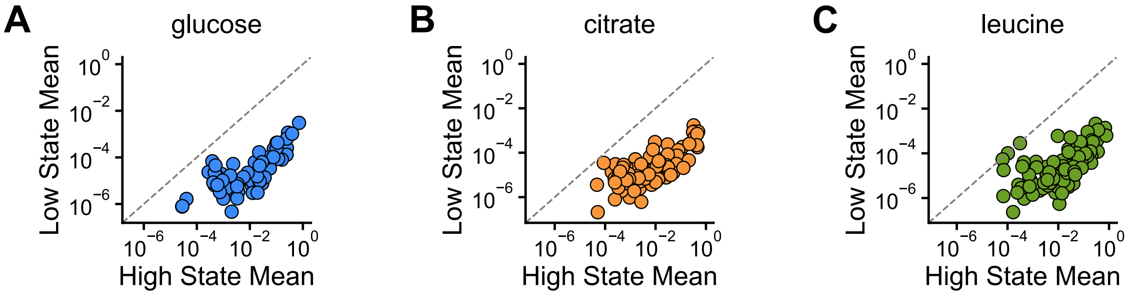


**Figure SI.12: Each ASV exhibits unique low- and high-abundance state means across multiple inocula experiments.** Scatter plots comparing mean relative abundances of low-abundance states (y-axis) vs high-abundance states (x-axis) for individual ASVs in (A) glucose, (B) citrate, and (C) leucine datasets. Dashed line indicates 1:1 line. Each point represents one ASV, demonstrating that abundance states are ASV-specific.

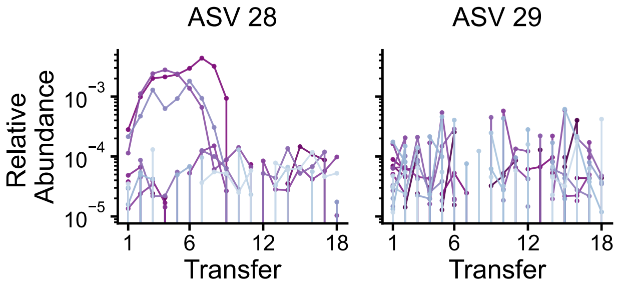


**Figure SI.13: Rare ASVs show consistent detection across transfers, ruling out contamination.** Time series of relative abundances for ASVs 28 and 29 across 18 transfers. Each colored line represents one of 20 replicate communities. Consistent detection throughout the experiment demonstrates these are genuine community members rather than contaminants.

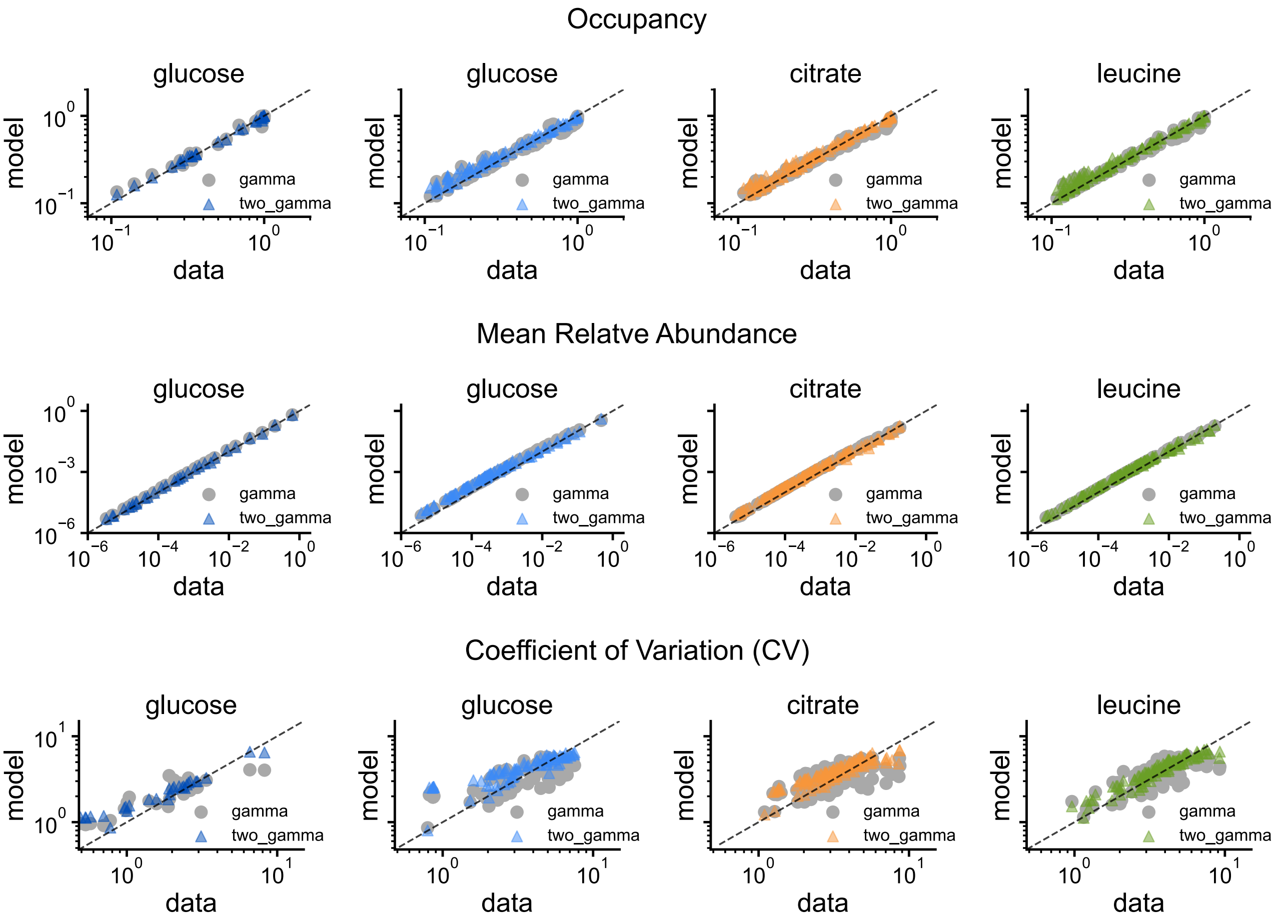


**Figure SI.14: Two-gamma model provides better predictions than single gamma model.** Comparison of empirical vs predicted values for (A) mean relative abundance, (B) occupancy, and (C) coefficient of variation (CV) for all dataset. While both models perform similarly for mean abundance and occupancy, the two-gamma model significantly outperforms the single gamma model in predicting coefficient of variation.

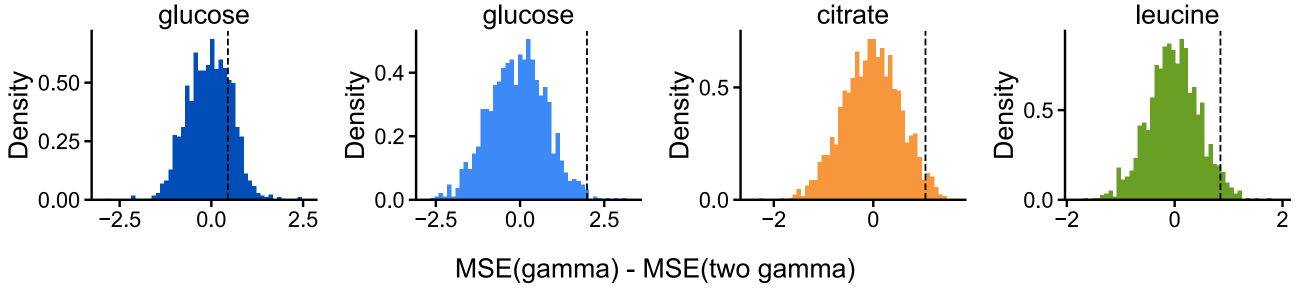


**Figure SI.15: Two-gamma model significantly outperforms gamma model in CV prediction.** Null distributions of MSE differences between models for coefficient of variation prediction. Empirical differences (dashed lines) show significantly better performance by the two-gamma model, particularly for multiple inocula datasets.

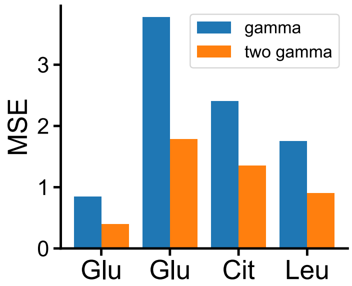


**Figure SI.16: Two-gamma model achieves approximately half the CV prediction error of gamma model.** Mean squared error (MSE) comparison between gamma and two-gamma models for coefficient of variation prediction across all datasets. The two-gamma model consistently shows lower MSE values, with approximately 50% less in prediction error compared to the single gamma model across all dataset.

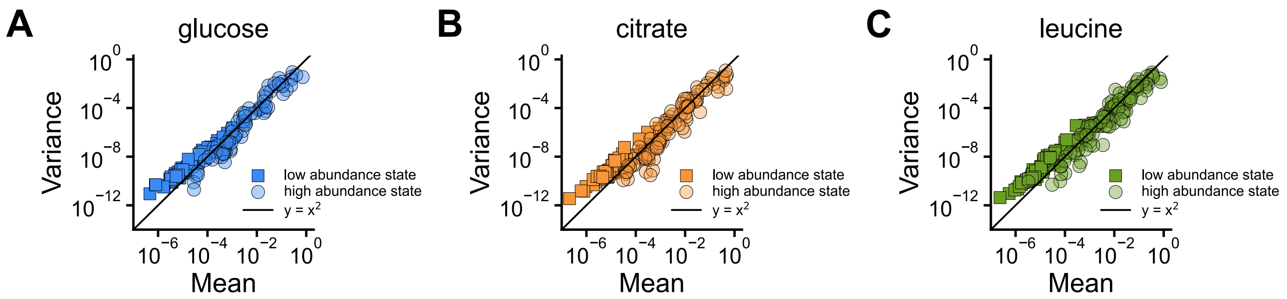


**Figure SI.17: Taylor's Law with exponent 2 holds for each state across all ASVs for the multiple inocula datasets.** Each panel shows Taylor's law for: (A) glucose, (B) citrate, (C) leucine.

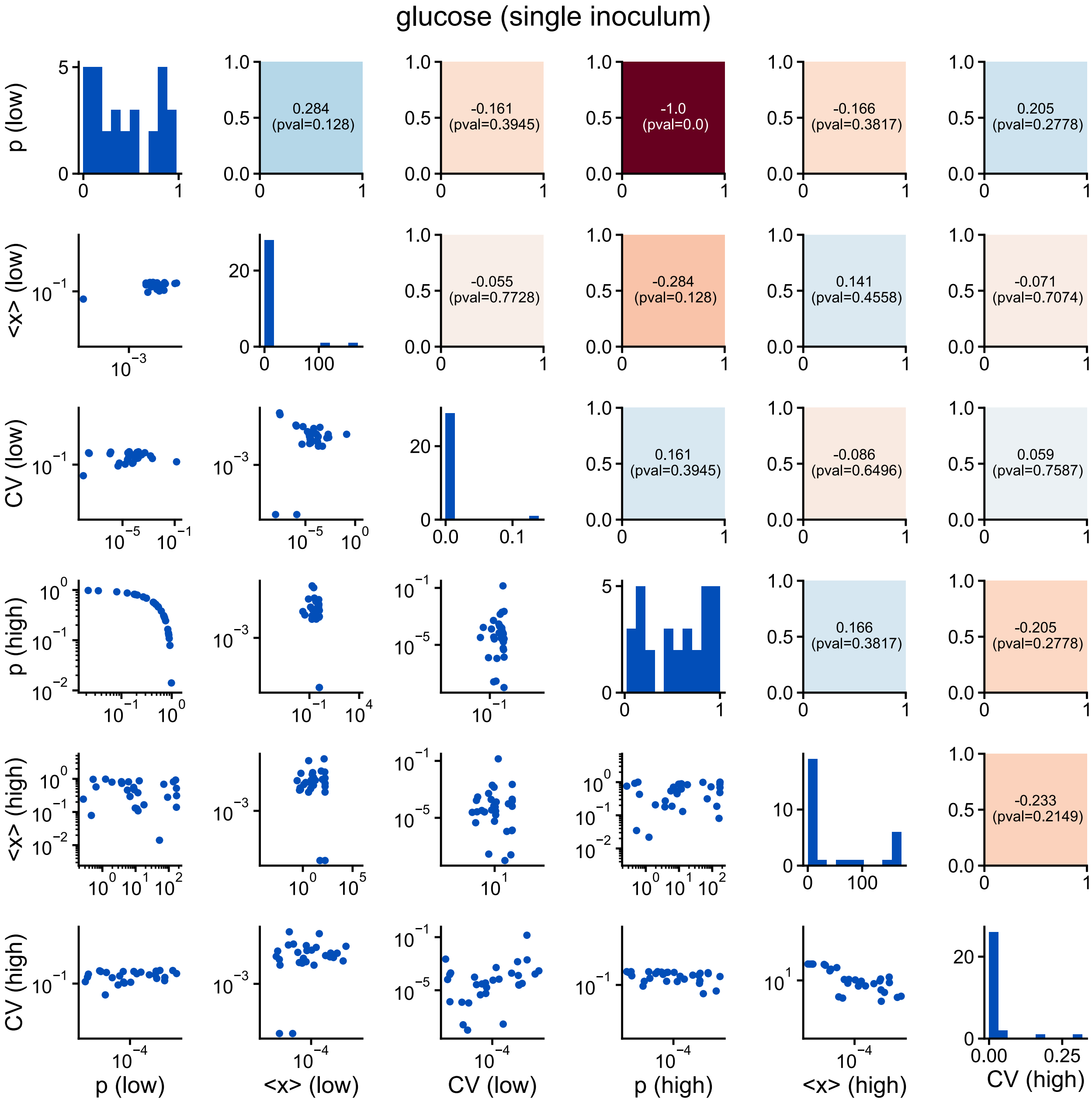


**Figure SI.18: Fitted gamma mixture parameters show weak correlations across ASVs for glucose (single inoculum).** Pairwise relationships among the five fitted gamma mixture parameters for glucose. Diagonal panels show parameter distributions, upper panels show correlation coefficients with p-values (colored backgrounds indicate significance), and lower panels show scatter plots. Most parameter pairs exhibit weak or non-significant correlations, demonstrating that the fitted mixture model parameters are largely independent of each other.

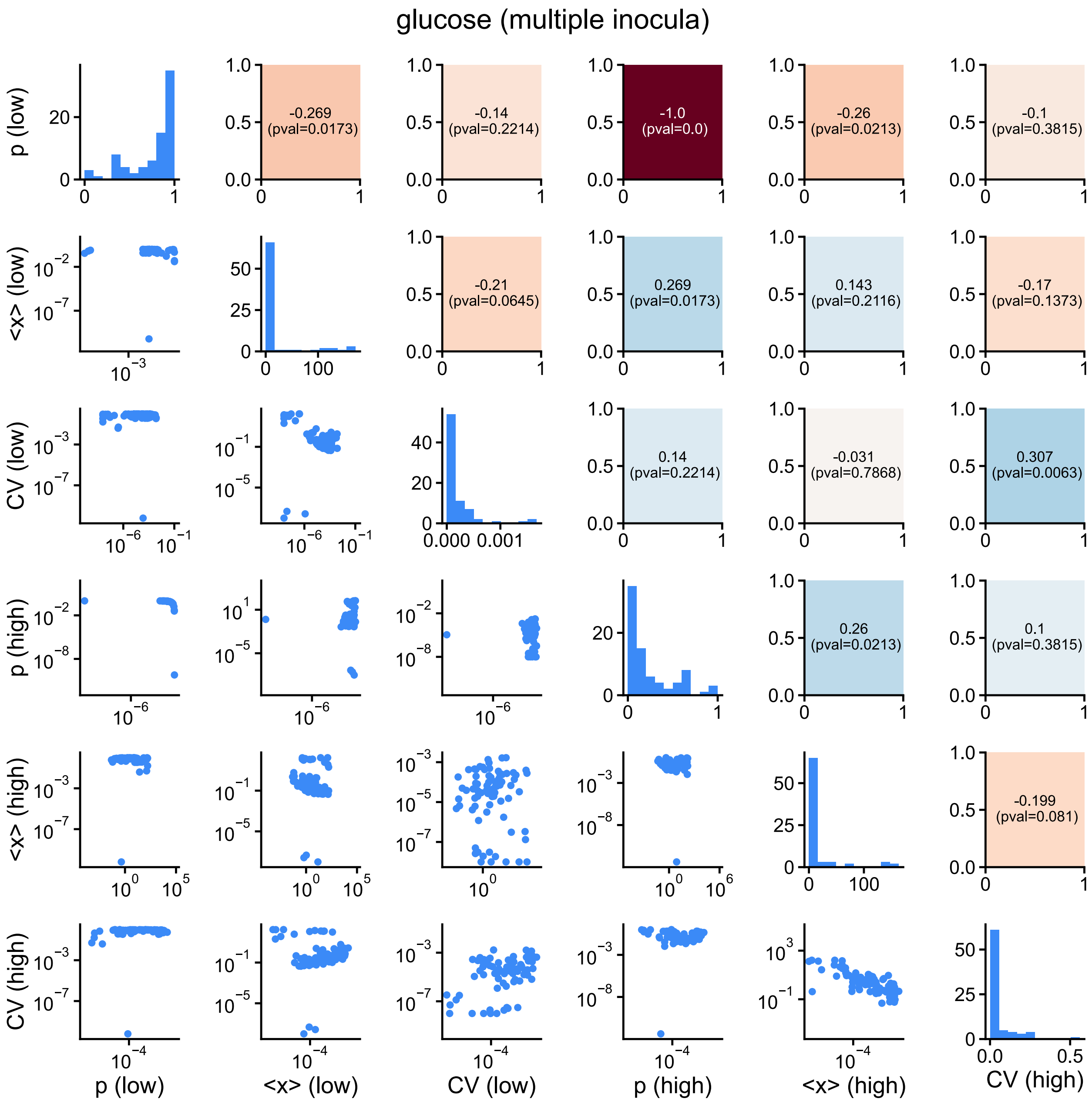


**Figure SI.19: Fitted gamma mixture parameters show weak correlations across ASVs for glucose (multiple inocula).** Pairwise relationships among the five fitted gamma mixture parameters for glucose. Diagonal panels show parameter distributions, upper panels show correlation coefficients with p-values (colored backgrounds indicate significance), and lower panels show scatter plots. Most parameter pairs exhibit weak or non-significant correlations, demonstrating that the fitted mixture model parameters are largely independent of each other.

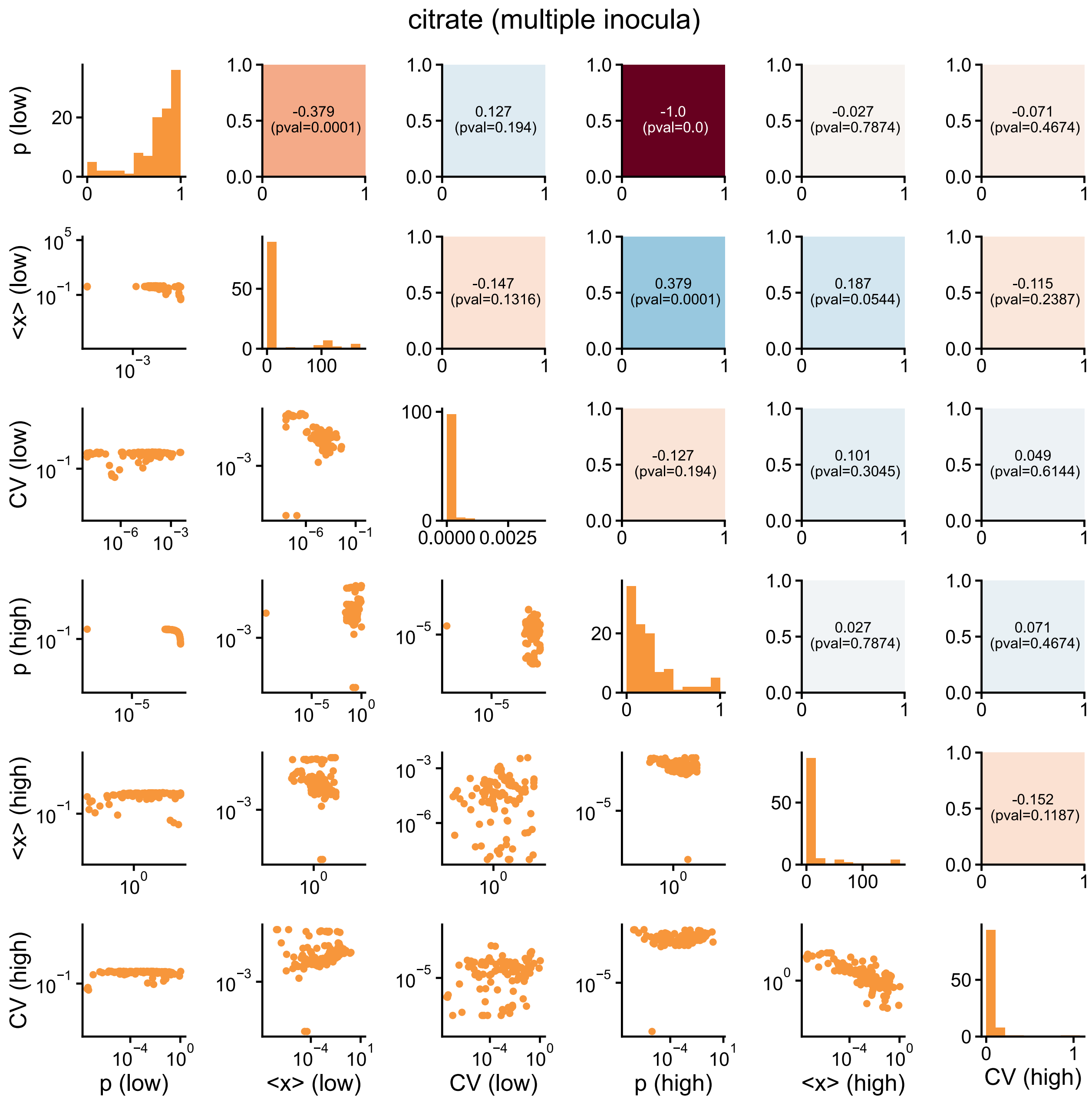


**Figure SI.20: Fitted gamma mixture parameters show weak correlations across ASVs for citrate (multiple inocula).** Pairwise relationships among the five fitted gamma mixture parameters for citrate. Diagonal panels show parameter distributions, upper panels show correlation coefficients with p-values (colored backgrounds indicate significance), and lower panels show scatter plots. Most parameter pairs exhibit weak or non-significant correlations, demonstrating that the fitted mixture model parameters are largely independent of each other.

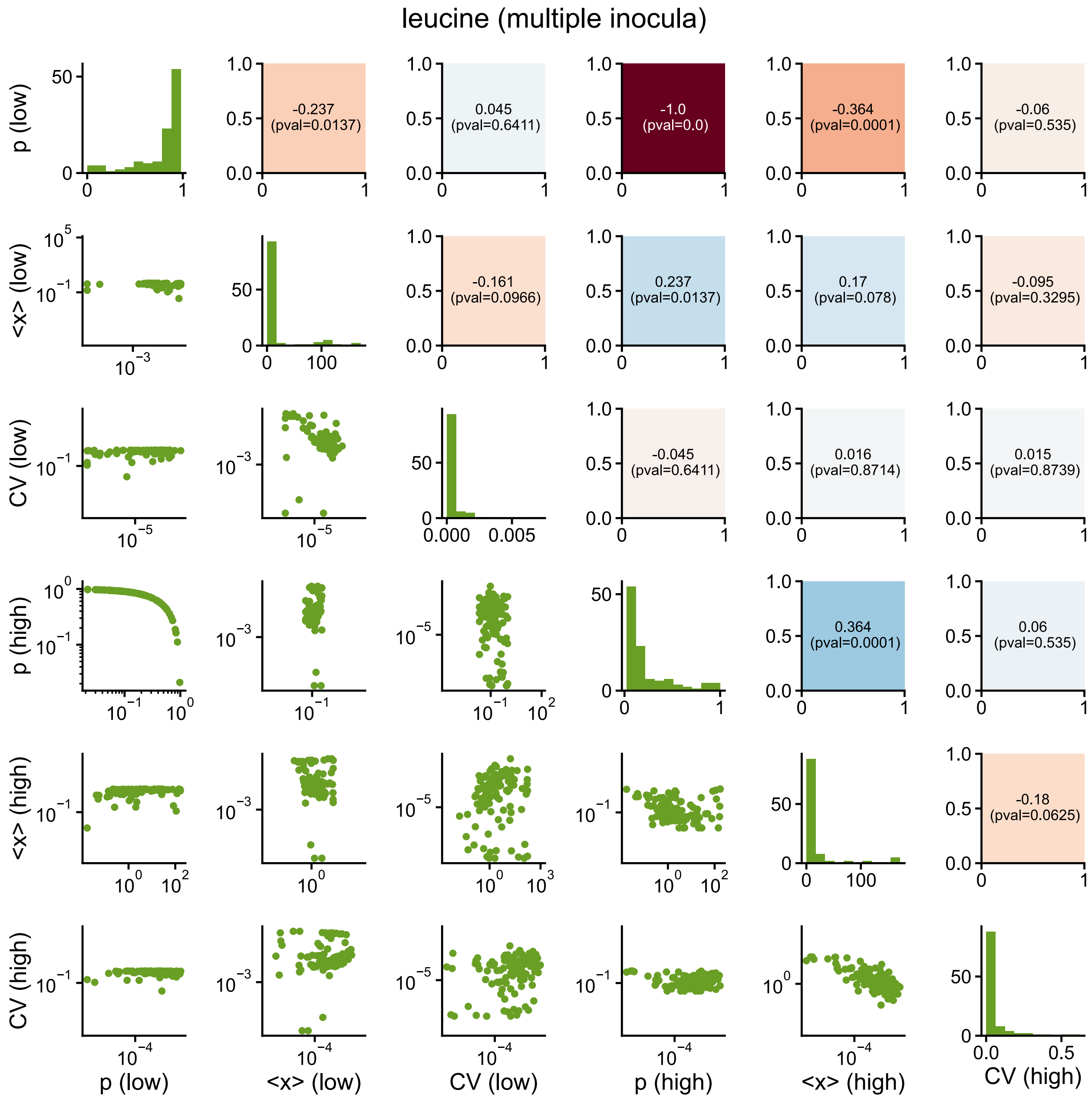


**Figure SI.21: Fitted gamma mixture parameters show weak correlations across ASVs for leucine (multiple inocula).** Pairwise relationships among the five fitted gamma mixture parameters for leucine. Diagonal panels show parameter distributions, upper panels show correlation coefficients with p-values (colored backgrounds indicate significance), and lower panels show scatter plots. Most parameter pairs exhibit weak or non-significant correlations, demonstrating that the fitted mixture model parameters are largely independent of each other.

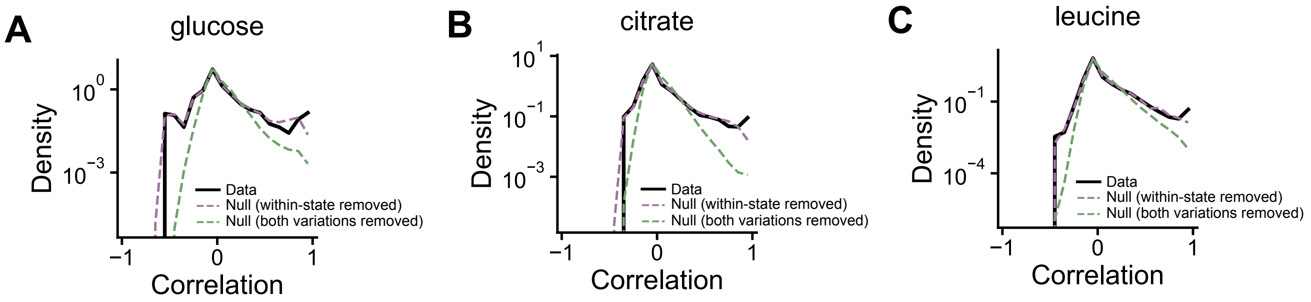


**Figure SI.22: Between-state variation drives abundance correlations across ASVs for the multiple inocula datasets.** The empirical distribution of pairwise ASV abundance correlations (black solid line) is compared to two null models for the three carbon sources of multiple inocula datasets: (A) glucose, (B) citrate, (C) leucine. A null model that removes within-state variation accurately reproduces the empirical correlations (violet dashed line). Meanwhile, a null model that removes between-state variation fails to reproduce the empirical correlations (green dashed line).

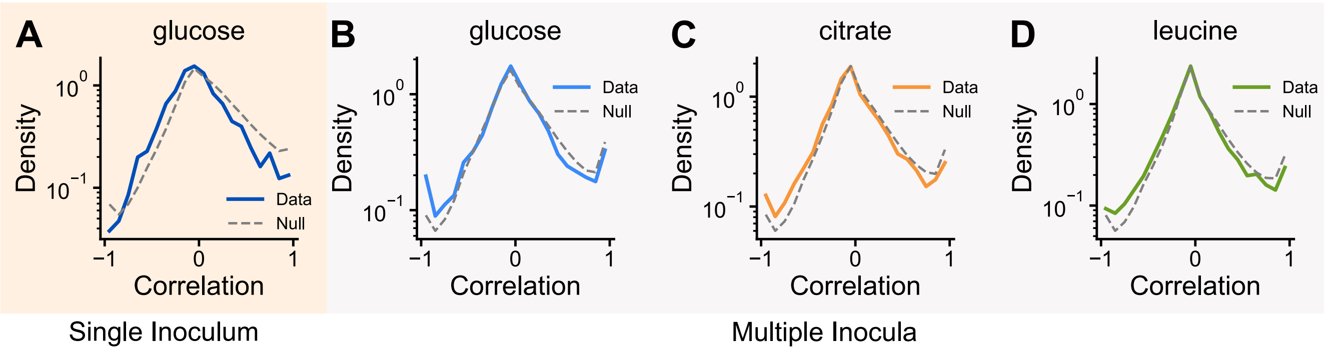


**Figure SI.23: Within-state variation does not contribute substantially to correlation structure across ASVs.** Distributions of pairwise ASV abundance correlations conditioned on states (solid lines) for single inoculum (A) and multiple inocula experiments: (B) glucose, (C) citrate, and (D) leucine. Correlations are shown for all state combinations: low-abundance and low-abundance states, high-abundance and high-abundance states, low-abundance and high-abundance states, and high-abundance and low-abundance states. Empirical distributions are compared to null model generated from in silico datasets by drawing abundances from fitted mixture of two gamma distributions (dashed lines).

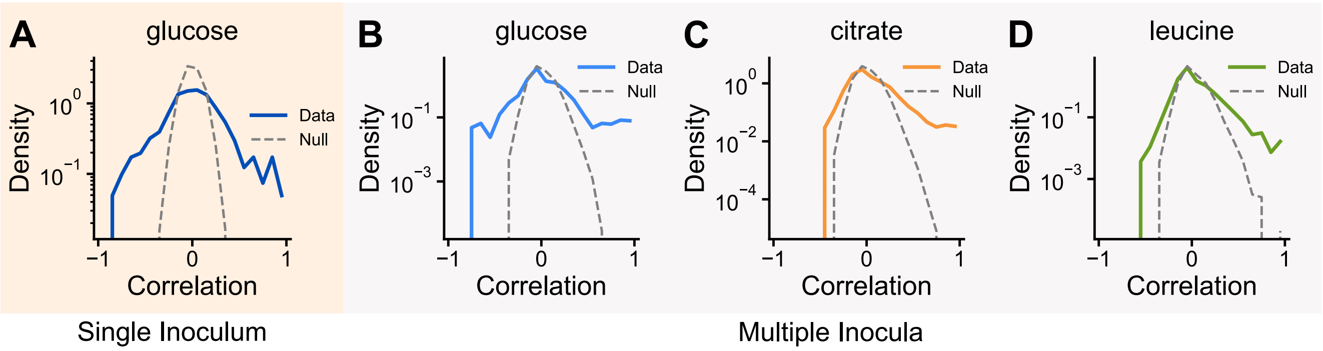


**Figure SI.24: Between-state variation strongly drives the correlation between ASVs.** Distributions of pairwise ASV state correlations ($\sigma$) (solid lines) for single inoculum (A) and multiple inocula experiments: (B) glucose, (C) citrate, and (D) leucine. Empirical distributions are compared to null model obtained by shuffling the variable $\sigma$ across communities (dashed lines).

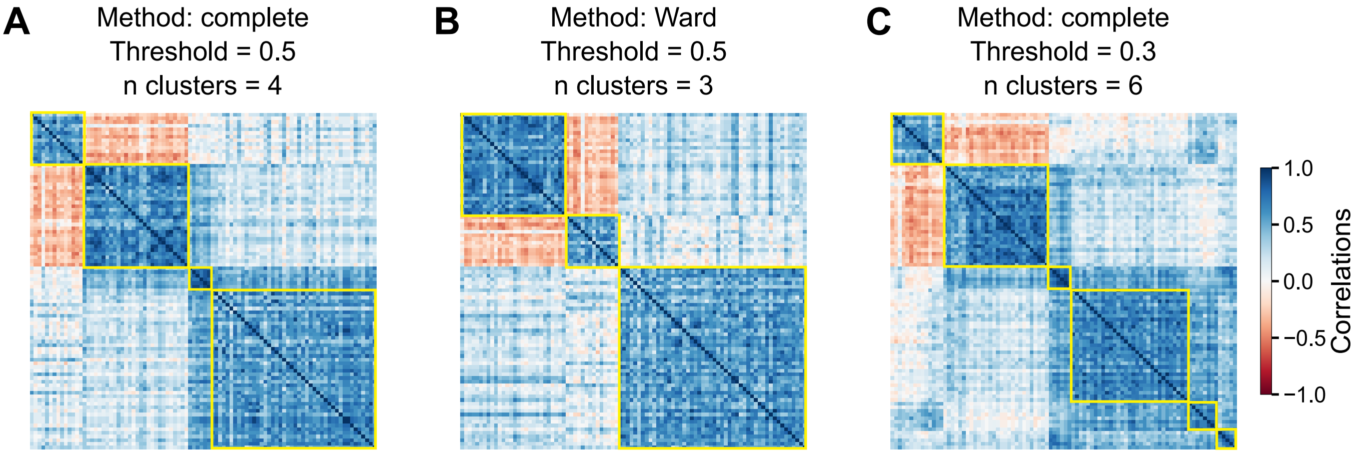


**Figure SI.25: Block structure in community correlations is robust to clustering method and threshold choice.** Community correlation matrices for the single-inoculum dataset are ordered using (A) complete linkage with threshold = 0.5 (4 clusters; shown in main Figure 4B), (B) Ward linkage with threshold = 0.5 (3 clusters), and (C) complete linkage with threshold = 0.3 (6 clusters). All approaches reveal clear block structure, indicating the presence of alternative stable community states. Complete linkage with threshold = 0.5 was used for all subsequent analyses.

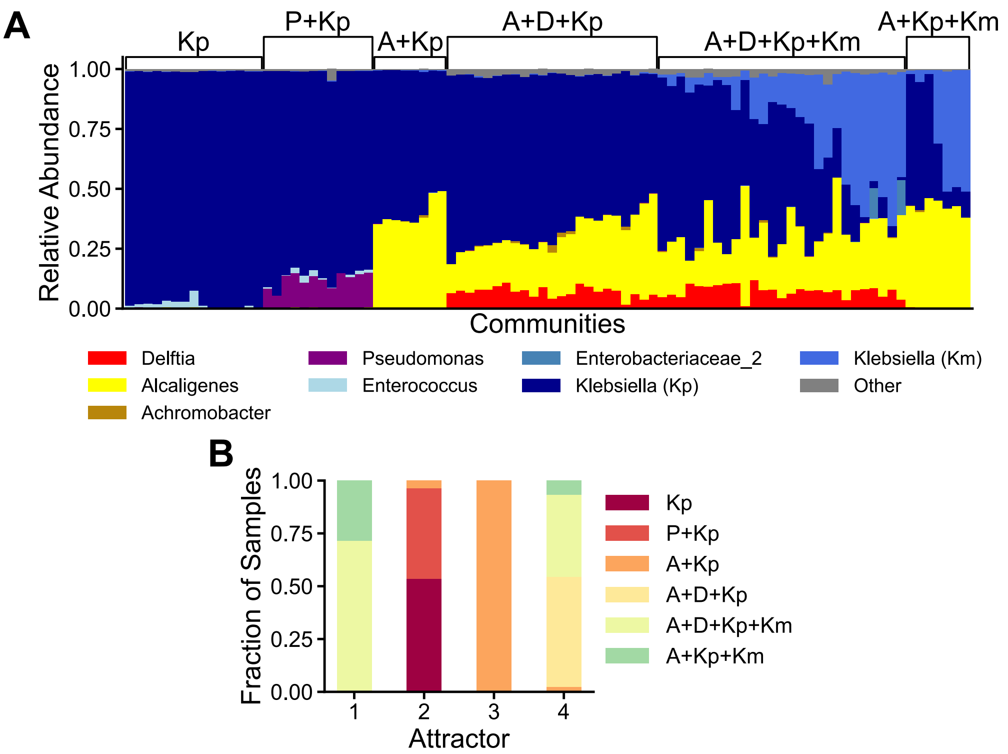


**Figure SI.26: Alternative stable states show non-trivial correspondence with taxonomic and functional compositions.** (A) Taxonomic composition at the ASV level for communities grouped by dominant taxa from Ref. [? ]. Each colored bar represents an ASV with relative abundance > 0.01; ASVs < 0.01 are grouped as Other. Labels indicate the six compositional groups based on dominant ASVs: Kp (*Klebsiella (Kp)*), P+Kp (*Pseudomonas* + *Klebsiella (Kp)*), A+Kp (*Alcaligenes* + *Klebsiella (Kp)*), A+D+Kp (*Alcaligenes* + *Delftia* + *Klebsiella (Kp)*), A+D+Kp+Km (*Alcaligenes* + *Delftia* + *Klebsiella (Kp)* + *Klebsiella (Km)*), and A+Kp+Km (*Alcaligenes* + *Klebsiella (Kp)* + *Klebsiella (Km)*). ASVs are functionally classified as fermenters (Klebsiella species, Enterobacteriaceae˙2, Enterococcus) or respirators (Pseudomonas, Achromobacter, Delftia, Alcaligenes) [? ]. (B) Mapping of the four alternative stable states from Figure 4B to the six taxonomic composition groups. Alternative stable states do not align perfectly with compositional classification but capture consistent community groupings.

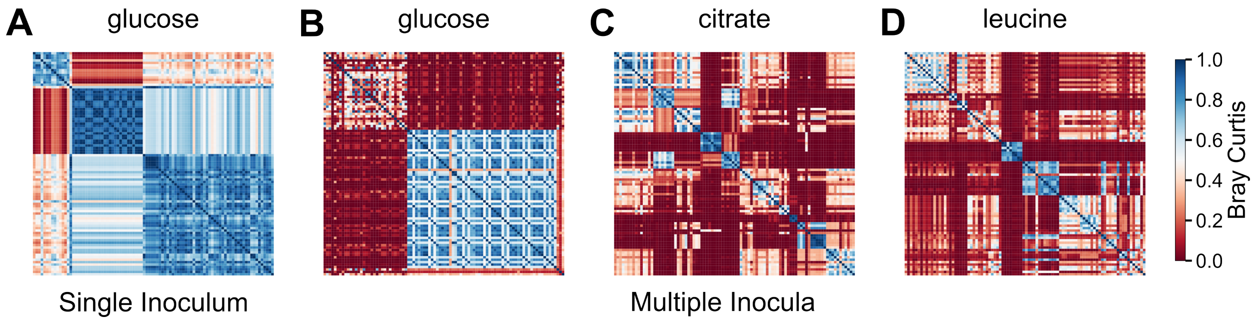


**Figure SI.27: Bray–Curtis dissimilarity reflects the structure of the correlation matrices.** Heatmaps display Bray–Curtis dissimilarity matrices in which samples are ordered according to the hierarchical clustering obtained from the corresponding correlation matrices. Panels show: (A) single inoculum (glucose; clustering from Figure 3B), and for multiple inocula: (B) glucose, (C) citrate, (D) leucine (clustering from Figure SI.29).

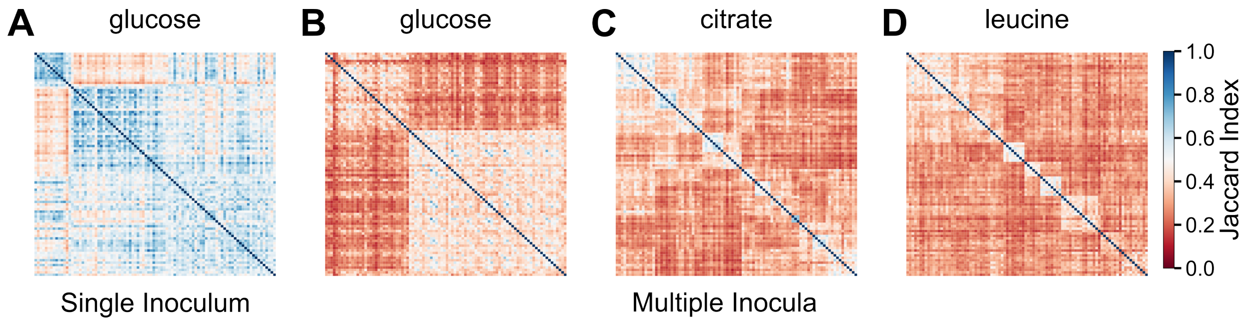


**Figure SI.28: Jaccard index also reflects the structure of the correlation matrices.** Heatmaps display Bray–Curtis dissimilarity matrices in which samples are ordered according to the hierarchical clustering obtained from the corresponding correlation matrices. Panels show: (A) single inoculum (glucose; clustering from Figure 3B), and for multiple inocula: (B) glucose, (C) citrate, (D) leucine (clustering from Figure SI.29).

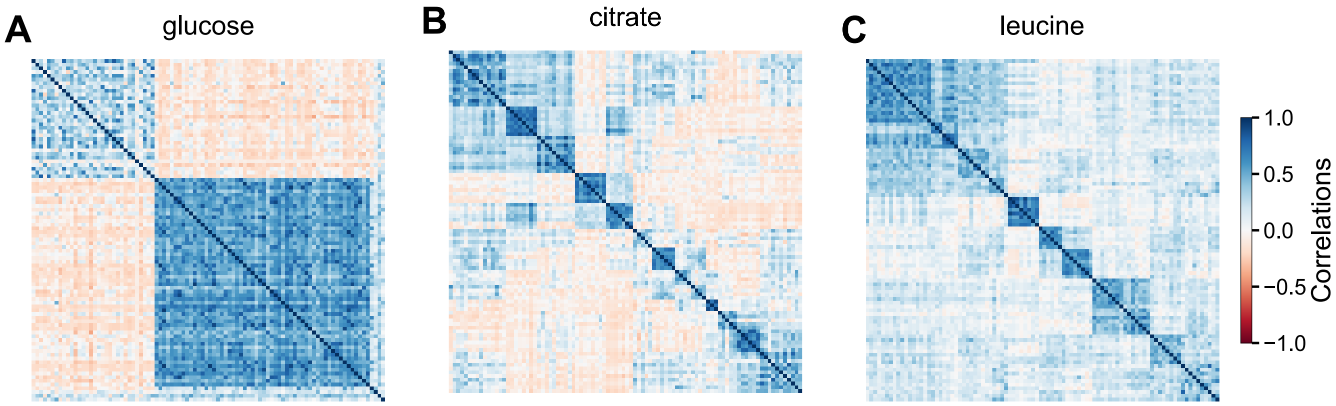


**Figure SI.29: Community correlation matrices for multiple inocula experiments also reveal block structures.** Correlation matrices based on binary state variables show distinct block structures after hierarchical clustering for (A) glucose, (B) citrate, and (C) leucine.

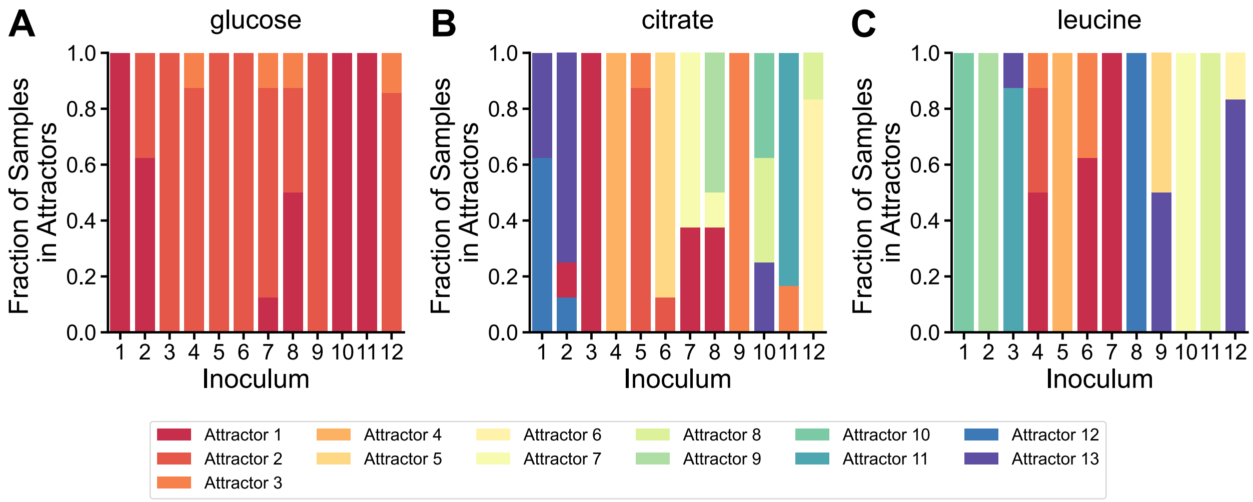


**Figure SI.30: Communities from the same inoculum tend to cluster in the same attractor.** Communities from each inoculum can be found in one or more alternative stable states for (A) glucose, (B) citrate, and (C) leucine. Each color represents a different alternative stable state. The predominance of single colors within each inoculum demonstrates that communities assembled from the same starting inoculum preferentially occupy the same alternative stable state.

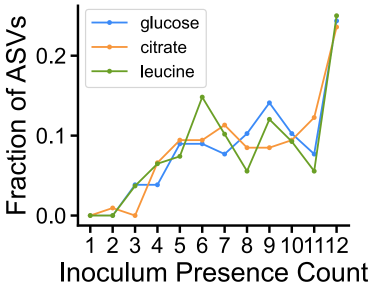


**Figure SI.31: ASV occurrence varies across inocula.** Fraction of ASVs present in different numbers of inocula for glucose, citrate, and leucine datasets. The distribution shows that many ASVs are detected only in subsets of inocula rather than universally.

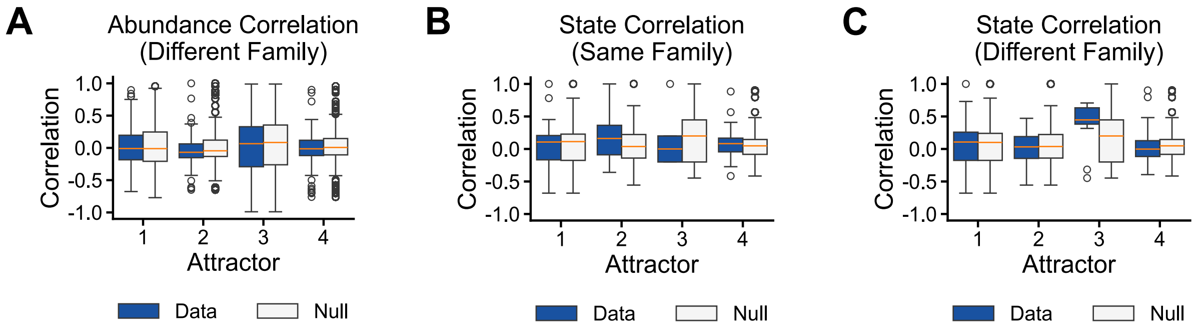


**Figure SI.32:** Within-attractor correlations show no family-level taxonomic signal. (A) Abundance correlations for ASV pairs from different families, (B) state correlations for ASV pairs within the same family, and (C) state correlations for ASV pairs from different families do not differ from a null model within each attractor.

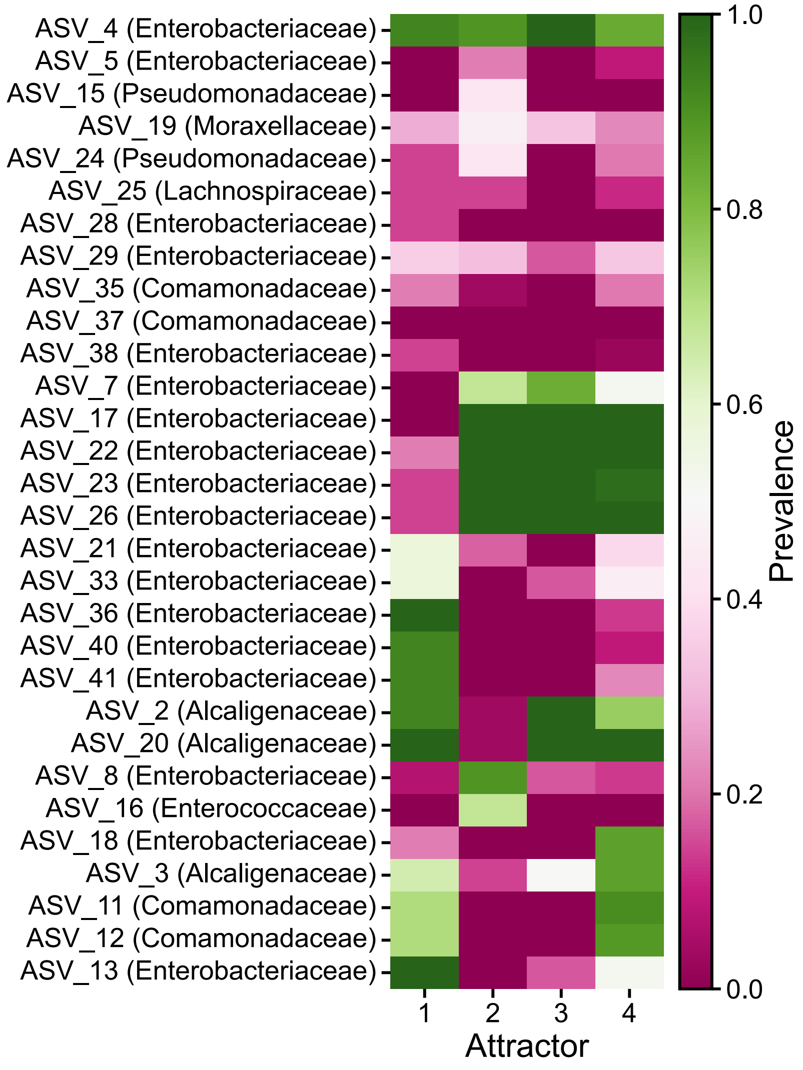


**Figure SI.33: ASV prevalence patterns with full identities and family information.** Heatmap of ASV prevalence across four attractors, corresponding to Figure 5A. Each row represents an ASV (with ASV name and family indicated), and each column represents an attractor. Colors indicate the probability of an ASV being in the high-abundance state.

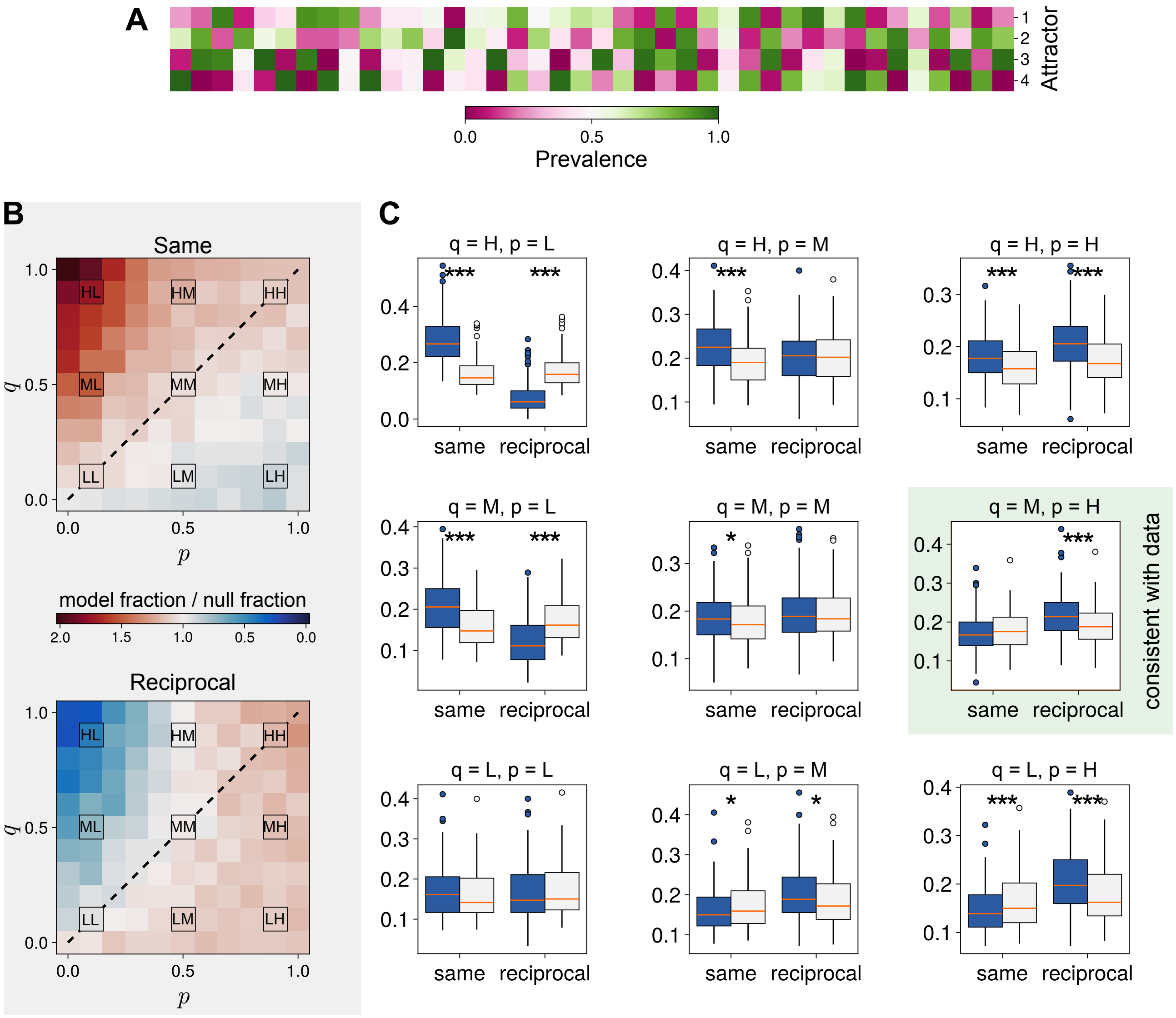


**Figure SI.34: Results of simulations of the Ising–Hopfield theoretical framework.** Simulations were performed with $N = 40$ ASVs partitioned into $F = 4$ equal-sized families, using $M = 4$ Hopfield memories. The overall exclusion strength was fixed to $J_{\text{mean}} = F - 1 = 3$, and dynamics were simulated at low stochasticity ($\beta = 100$) using $N_{\text{sweep}} = 1000$ Monte Carlo sweeps per run. For each parameter pair $(q, p)$, we generated $N_{\text{runs}} = 200$ independent assembly runs per realization of the memories, and averaged results over $K = 1$ (A), $K = 20$ (A) and $K = 200$ (B) independent memory realizations. We show (A) an example of a prevalence pattern for $q = 0.5$ and $p = 0.9$. We report (B) heatmaps showing enrichment ratios for within-family prevalence patterns as a function of the filtering parameter $q$ and exclusion contrast $p$. The color scale reports the ratio between the simulated fraction of ASV pairs under a null model obtained by shuffling family labels while preserving the prevalence-pattern matrix, for both *same* and *reciprocal* fraction. Values larger than one indicate enrichment relative to the null. The diagonal line indicates the analytical scaling $q = p$ predicted by the low-stochasticity mean-field analysis. We also report (C) boxplots comparing simulated fractions and matched-null expectations for representative parameter combinations of low (L), medium (M), and high (H) values, corresponding to 0.1, 0.5, and 0.9. For each case, we show distributions over $K = 200$ independent realizations of the model of (blue) simulated within-family fractions and (gray) the corresponding null means obtained by shuffling family assignments within the same prevalence patterns. Left boxes correspond to same-pattern fractions, right boxes to reciprocal-pattern fractions. Statistical significance is assessed using two-sided permutation tests, with $10^4$ random permutations per comparison, comparing simulated and null distributions across realizations; asterisks denote significant deviations from the null with $*$ ($< 0.05$), $**$ ($< 0.01$), $***$ ($< 0.001$). Only the combination $q = M, p = H$ is consistent with the data, indicating that both environmental filtering and limiting similarity play a non-negligible role in shaping community composition, with competitive exclusion due to limiting similarity being the dominant force.
